# Robust identification of cell-cell communication heterogeneity in single cells

**DOI:** 10.64898/2026.04.29.721691

**Authors:** Federico Bocci, Yunlong Jia, Scott Atwood, Qing Nie

## Abstract

Communication between cells modulates cell fate decisions by relaying information across tissues and inducing intracellular responses mediated by gene regulatory networks. Inference of cell-cell communication from high throughput data such as single cell transcriptomics is gaining popularity due to the high data availability and ease to automate modeling over hundreds of signaling pathways. Studying how cell-cell communication operates across biological scales and influences cell fate decisions, however, remain a major open question. Here, we present scRICH, a framework and package that integrates mechanism-based, multiscale mathematical modeling with learning strategies to capture the complexity of cell-cell communication from single-cell and spatial transcriptomics data. scRICH unravels the heterogeneity of communication behavior within cell types, links cell-cell communication to cell fate decisions by incorporating dynamical information of RNA splicing, and connects the scales of cell-cell interactions and intracellular response by constructing multilayer regulatory networks. We validate scRICH with new experiments on EGF ligand/receptor co-expression in keratinocytes from skin-equivalent organoid, and compare these computational predictions against existing CCC inference methods. Applying scRICH to multiple biological scenarios demonstrate its ability to capture emerging relations between distinct cell-cell communication pathways, interactions at the onset of cell fate decision, and emerging trends in cell-cell communications along cell lineages and in space.

## Introduction

Cell-cell communication (CCC) inference from single cell sequencing data has now gained popularity as a highly automated, hypothesis-free approach to discover new molecular interactions^1,2^. Despite the rapid progress in the field, several existing limitations hamper the interpretability and scope of these methodologies.

First, CCC inference strikes a delicate balance between high resolution, achieved with models at the scale of individual cells, and robustness, achieved by averaging gene expression over large groups of cells – typically the annotated cell types in a dataset. Several methods exploit the single cell resolution offered by state-of-the-art transcriptomics to predict CCC between individual cells, thus resulting in dense interaction networks^3–7^. A major drawback of this approach is the sparsity in single cell transcriptomics data^8^, which can result in large cell-to-cell fluctuations in gene counts and missing information for important signaling pathways. To ensure robust predictions, many popular methodologies leverage cell annotations to average the gene expression and/or the CCC connections over cell types^9–13^. The major pitfall of this approach is the potential loss of information about heterogeneity within individual cell types. In other words, sharing the same cell type does not necessarily imply a similar role in CCC, and cells of a given type can either send, receive, or do not participate to a given signaling pathway.

Second, it is currently difficult to inject dynamical information about cell fate specification and its mutual dependence with CCC. Despite the abundance of methodologies to infer lineages and cell transition from single cell data, cells are typically modeled in an autonomous manner^14,15^. A “static” view of CCC where cell states are treated as stable (i.e., not transitioning toward new cell states) and homogeneous (i.e., all cells behave similarly given a ligand/receptor pair) prevents a complete understanding of how CCC is modulated, and in turns contributes to cell fate decision events. For such approach to be possible, the cell states and “CCC propensity” need to be treated as distinct observations, whereby for example the former is defined by well-defined marker genes and morphological changes, whereas the latter is defined by ligand/receptor expression.

Third, cell-cell communication is multiscale as signal transduction networks “process” external information, modulating secondary CCC pathways in a constant back-and-forth interaction between the intercellular and intracellular scales. Recently, a few methods included information about receptor downstream targets, which can be used as a benchmark to confirm pathway activation^3,16–23^. For example, SoptSC refines the single cell-level CCC networks by weighting communication edges based on the expression of a downstream target gene set that is provided by the user^3^. Similarly, NicheNet and scMLnet feature manually curated databases to link ligand-receptor pairs to candidate downstream targets^16,17^, while CytoTalk identifies putative response genes *de novo* using a mutual information approach^18^, and LIANA+ integrates CCC inference with database knowledge^19^. While these methods took decisive steps forward toward connecting the ligand/receptor and downstream signaling levels, we propose that the recent advances in gene regulatory networks (GRN) reconstruction from single cell transcriptomics^24^ could now allow a more holistic and detailed description of the “cellular flow” that starts from ligand/receptor interactions, leads to a downstream response, and in turn regulates other CCC pathways.

Here, we introduce scRICH (<u>R</u>obust identification of <u>c</u>ell-cell communication <u>h</u>eterogeneity in single cells), a methodology that addresses these existing bottlenecks by integrating cell-cell communication, cell-fate transitions directionality via RNA velocity, and gene regulatory networks (GRN) inference to achieve a multiscale representation of cell-cell communication. scRICH takes scRNA-seq counts and well-annotated cell labels as input to (1) identify robust CCC modes across cell types; (2) integrate downstream target RNA velocity into a multilayered model of cell-cell signaling; (3) enable the simultaneous visualization of transition trajectories and CCC networks; and (4) connect ligands-receptor interactions to their downstream gene regulatory network (GRN) to predict the connection between distinct cell-cell communication pathways. We apply scRICH to multiple biological scenarios including keratinocyte differentiation, erythroid cell differentiation, and cancer epithelial-mesenchymal transition. Specifically, we confirm experimentally scRICH’s prediction about EGF signaling heterogeneity and relation with other CCC pathways including TGFb, NOTCH, and CDH4. Finally, we demonstrate scRICH’s ability to identify the spatial localization of cells with coherent CCC behavior in heterogenous tissues via analysis of spatial transcriptomics (ST) data.

## Results

### Overview of scRICH

scRICH takes the preprocessed, log-normalized scRNA-seq counts and well-annotated cell labels as input (Fig **1A**, top). While “regular” scRNA-seq counts can be used, some of the downstream analysis tools explicitly requires the unspliced and spliced count matrices for CCC validation and projection onto cell trajectories.

**Figure 1.**
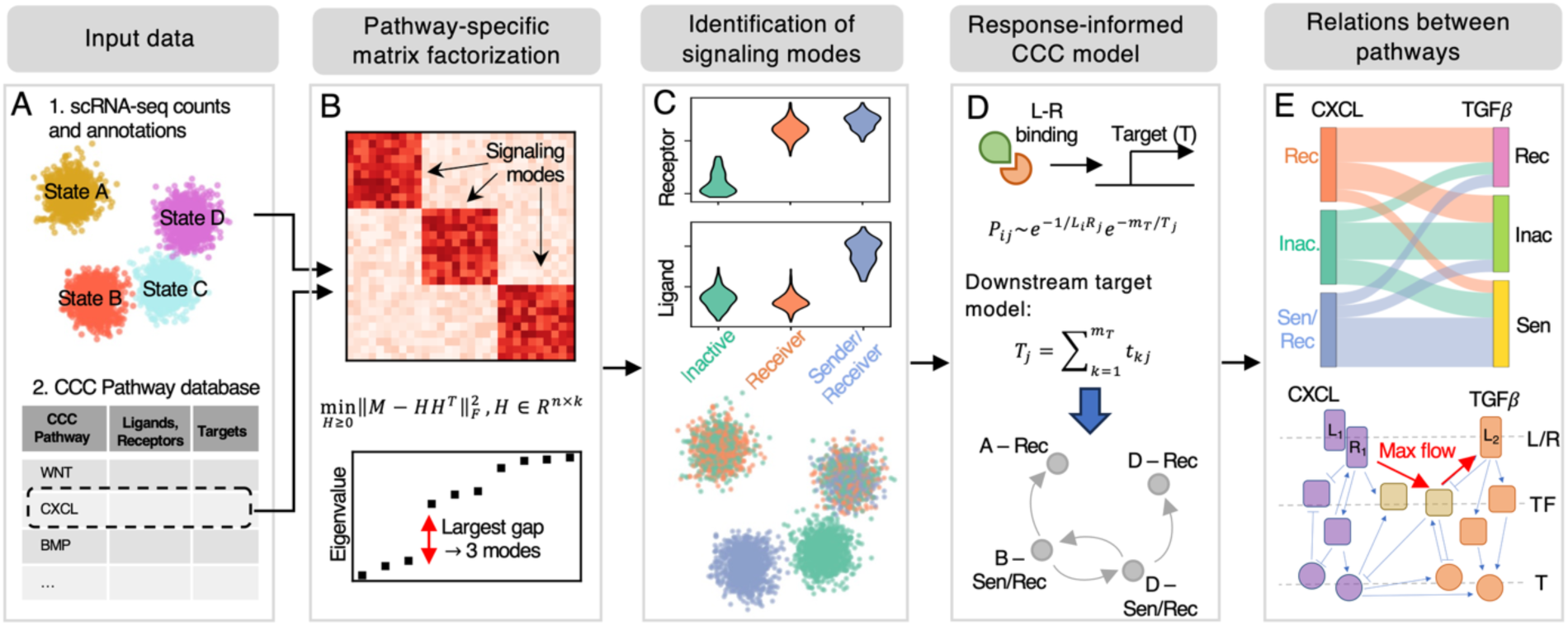
Overview of scRICH. **(A)** The inputs to scRICH are the well-annotated scRNA-seq counts and database information on pathways ligand/receptor, transcription factors (TF) and downstream targets (T). **(B)** For each CCC pathway, scRICH construct a cell-cell similarity matrix. The optimal number of CCC modes is computed via nonnegative matrix factorization and spectral analysis. **(C)** The CCC signaling modes are annotated based on expression of receptors and ligands, and their correspondence to cell types is quantified. **(D)** A CCC model is constructed whereby the propensity of communication depends on ligand and receptor abundance in the senders and receivers as well as downstream target activation, which is quantified either by gene expression or RNA velocity. **(E)** The relationship between CCC signaling modes of distinct pathways is identified, and the molecular connection between pathways is discovered by constructing the joint GRN. Finally, a max flow strategy identifies the regulatory connections in the GRN.

### Quantification of intra-type cell-cell communication heterogeneity

First, given a specific CCC pathway, scRICH dissects the CCC heterogeneity with unsupervised clustering based exclusively on the transcriptomics counts of ligands, receptors and downstream targets (if known) of the pathway (Fig **1A**, bottom). To compile a comprehensive gene set for each CCC pathway, scRICH assembles ligands/receptors sets from CellChat^9^ and downstream targets from exFinder^25^, which is in turn assembled based on several existing databases^16,26,27^ (Supplementary Fig. **S1A-H**). Alternatively, pathway gene signatures compiled from the gene set enrichment analysis (GSEA) molecular signature database^28^ or custom-made gene set can be provided as input. scRICH determines the optimal number of CCC modes in the dataset with nonnegative matrix factorization of the pathway-specific cell-cell similarity matrix (Fig. **1B**), an approach that has been previously employed to identify cell states in single cell transcriptomics data^29^. For clarity, the terms “cell state” and “cell type” will indicate the cell annotation based on marker analysis, which is provided as input to scRICH, whereas the term “mode” will indicate the pathway-specific CCC clusters identified by scRICH. To achieve robust, sub-type resolution, the cell types and CCC modes are integrated in meta-cells, whereby a dataset with *N* cell types and *M* signaling modes will be divided into *NxM* meta-cells, with the distinctive advantage that cells within each meta-cell exhibit consistent transcriptomics profile and similar CCC behavior (Fig. **1C**). A heterogeneity score is computed to quantify whether cells of same type have consistent CCC behavior (low score) or exhibit heterogenous CCC behaviors (high score). This pathway specific clustering is repeated in parallel for all pathways in the scRICH database, whereby user-defined parameters can filter the number of analyzed pathways. Finally, a statistical test based on permutation is implemented at a pathway-specific level to quantify the merit of splitting cell types into meta-cells for exploration of CCC behavior heterogeneity and downstream analysis.

### Construction of communication networks and transitions

Starting from the CCC meta-cells defined in the previous step, scRICH builds a model of CCC between meta-cells through the given pathway (Fig **1D**). In the model, the signaling between two meta-cells is a function of ligand and receptor expression in the sender and receiver meta-cells, respectively, as well as the activation of downstream targets in the receiver meta-cell. Specifically, scRICH offers two mathematical models of the ligand-receptor interaction. First, a “diffusion-based” model where the communication propensity depends exponentially on the expression of ligands and receptors, similar to the CCC model at individual cell scale employed by SoptSC^3^; in alternative, a “mass action-based” model similar to CellChat whereby the communication propensity depends on ligand and receptor expression via a Michaelis-Menten-like model^9^. While both models provide the same qualitative output, the diffusion-based model amplifies strongest interactions, and could thus be a better choice to simplify complex systems with several cell states and CCC signaling modes. Conversely, the mass action-based model is more permissive and highlights weaker interactions between meta-cells, perhaps being better suited to study smaller systems (Supplementary Fig. **S1I-L**). The activation of downstream targets is quantified by the target RNA velocity, which indicates whether the gene is being upregulated or downregulated, respectively^30,31^. Alternatively, “regular” RNA counts are used if unspliced/spliced counts are not provided in the input dataset. The inclusion of downstream targets is optional, and the method can be run using exclusively the information on receptors and ligands. The heterogeneity of cell-cell communication within a cell type can be visualized in low-dimensional embedding by color-coding cells based on CCC signaling mode membership (Fig. **1C**, bottom).

### Cellular flow between co-dependent pathways

Finally, scRICH integrates the information on cell clustering based on individual CCC pathways to extrapolate emerging biological trends and predict the “cellular flow” between CCC pathways (Fig **1E**). First, scRICH uses mutual information to identify CCC pathways that show similar trends across cell states (for example, being a WNT sender implies being a BMP sender, etc…) and employs Sankey-style diagrams to represent the overlap between signaling modes of distinct CCC pathways. Moreover, given two distinct CCC pathways, scRICH implements GRN inference using the spliceJAC package^32^ to reconstruct the joint regulatory network including the ligands, receptors, and downstream transcription factors connecting the two pathways, and uses a maximal flow algorithm to identify the regulatory connections between two distinct pathways.

### Benchmarking and comparison with other methods

We benchmark scRICH’s ability to predict cell-cell communication networks by extensively comparing against existing tools for CCC inference including CellChat^9^, CellPhoneDB^11^, and Liana+^19^. First, we demonstrate similar accuracy when scRICH is used to infer CCC at the cell type level, defined as the pre-existing cell annotation in the dataset. Second, we highlight novel insights uniquely provided by scRICH by reconstructing the CCC at meta-cell resolution in our own single-cell RNA-seq dataset for fibroblast-containing human skin equivalents^33^ (FibHSEs). We start by identifying the top CCC pathways based on the average expression of receptors and ligands using scRICH’s built-in CCC pathway exploration function (Fig. **2A**). For a given CCC pathway, scRICH first identifies the number of CCC communication modes. In the example of the EGF pathway, scRICH identifies three signaling modes corresponding to EGF inactive (low expression of receptors and ligands), EGF receiver (high receptor, low ligand), and EGF sender/receiver (high receptor, high ligand). These EGF CCC modes can be visualized in low-dimensional embedding as well as via violin plots (Fig. **2B-C**). To mitigate sparsity in the signaling mode identification, scRICH utilizes imputed counts for genes associated to ligands, receptors, and targets, potentially introducing a dependence on imputation parameters including number of principal components (PCs) and cell neighbors (N). We evaluated this dependence by computing the Wassertein distance between signaling mode memberships obtained for different (PC, N) parameter combinations for the top 10 identified pathways in the FibHSE dataset. While the number of PCs does not influence signaling mode identification, cell membership to different signaling modes depends, albeit weakly, to the number of cell neighbors (Fig. **S2A**). Specifically, some pathways including Annexin, APP, IGF, and NRG exhibit different numbers of signaling modes for varying parameter combinations (Fig. **S2B**), with occurrences of signaling modes splitting or merging (Fig. **S2C-F**). Repeating this analysis on other datasets with varying number of cells and cell types confirmed the robustness with respect to number of principal components and cell neighbors (Fig. **S2G-J**). Afterwards, scRICH constructs the EGF communication network detailing how the inactive, receiver and sender/receiver subsets of the epidermal cells communicate through EGF (Fig. **2D-E**). Given that CellChat, CellPhoneDB, and Liana+ infer communication networks at the cell type level, we coarse-grain scRICH’s prediction by averaging over CCC modes, thus predicting a CCC network that can be directly compared (Fig. **2F-G**). We repeat this procedure over several CCC pathways presented in Fig. 2A and compare the predicted CCC matrix based on (1) spearman correlation, (2) element-wise matrix distance, and (3) overlap of predicted interactions. Overall, we found strong overlap between ScRICH and the CCC networks predicted by CellChat, CellPhoneDB, and Liana+ when applying both the mass-action and diffusion models of cell-cell communication (Fig. **2H**). scRICH can estimate CCC interactions based on (1) standalone ligand/receptor expression; (2) ligand/receptor and target gene expression; or (3) ligand/receptor expression plus RNA velocity of target genes. Testing these different CCC models across three datasets suggests that the relation between these models could be dataset-specific, likely due to context-specificity in target gene selection (Fig. **S2K-M**). Furthermore, we compared different RNA velocity inference models, scVelo and Unitvelo, finding that scVelo-based prediction agrees closely to target-free and target expression-based models (Fig. **S2M**). These results highlight the important context-specific target gene selection and RNA velocity model – both customizable options – when applying scRICH.

**Figure 2.**
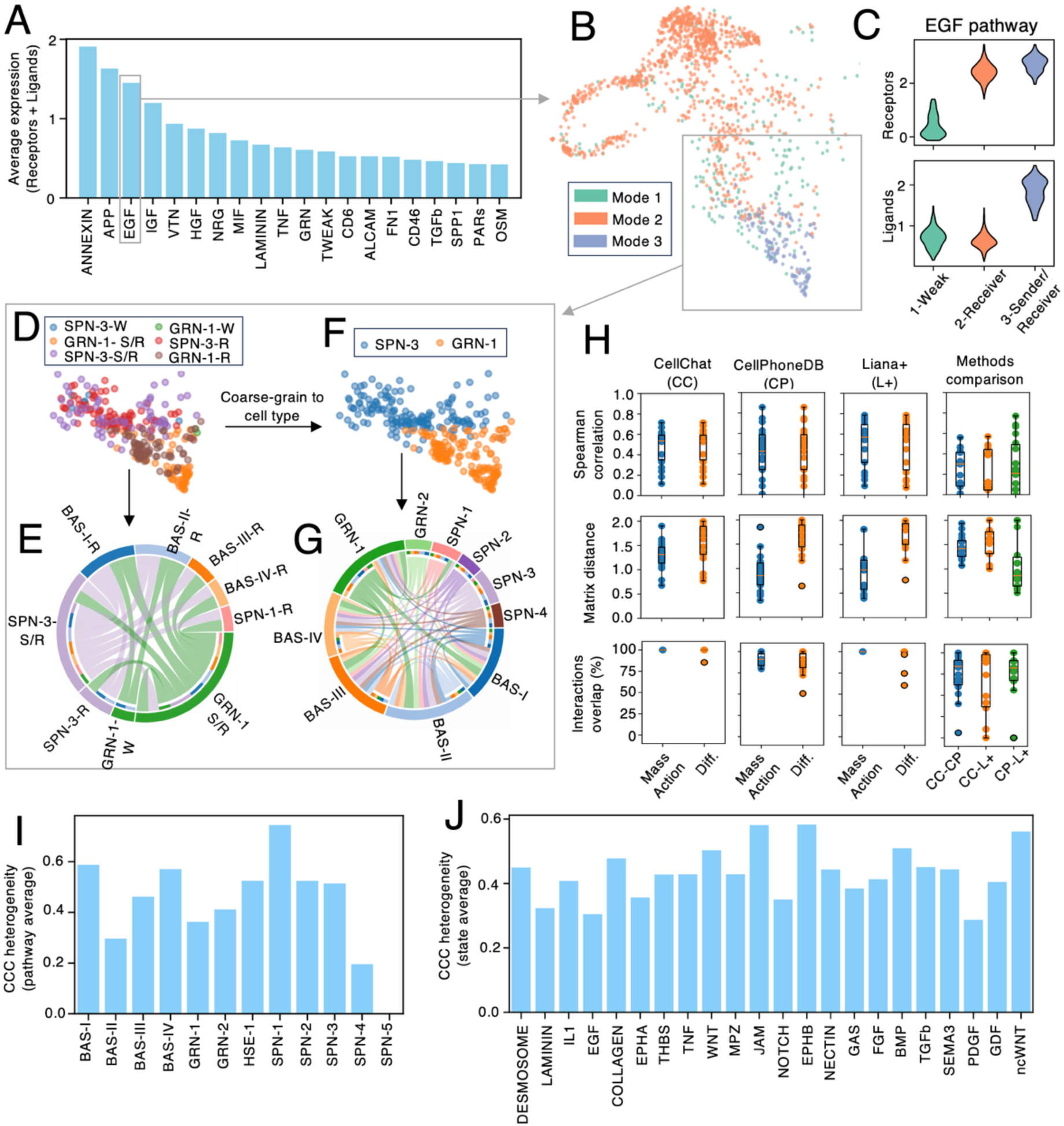
Comparison of scRICH against existing CCC inference methods. **(A)** The top CCC signaling pathways identified by scRICH in FibHSE organoids based on average expression of ligands and receptors. **(B)** Low-dimensional embedding whereby FibHSE cells are color-coded with EGF CCC communication modes. **(C)** The expression (normalized and log-transformed) of EGF receptors and ligands in the identified communication modes corresponding to an inactive (green), receiver (orange), and sender/receiver (blue) states. **(D-G)** Schematic of the quantification pipeline. (D) The UMAP at sub-cluster resolution showing how epidermal cell states split at the meta-cell level based on the EGF communication modes. (E) The EGF CCC network predicted by ScRICH at meta-cell resolution. (F) Cell state level UMAP representation obtained by aggregating meta-cells over epidermal cell states. (G) The EGF CCC network at cell type resolution. **(H)** Detailed comparison between ScRICH and the CCC networks predicted by CellChat, CellPhoneDB, and Liana+. The rightmost column shows the comparison between the three methods as benchmark. The line, box and whiskers depict average, first quartile (Q1) to third quartile (Q3) range, and furthest point within 1.5x the interquartile range, respectively. **(I)** The cell type CCC heterogeneity when aggregating results from all considered CCC pathways. **(J)** The CCC signaling pathway heterogeneity when aggregating over all cell types.

Finally, we focus on the information loss resulting from averaging CCC networks at the cell type level by defining and computing a CCC heterogeneity score bound within [0,1]. This score quantifies whether cells with different roles in the CCC pathway (senders, receivers, *etc.*) are uniformly distributed across cell types (high score) or clearly divided along cell types (low score). Several keratinocyte states such as BAS-I and SPN-1 exhibit a high score, i.e., cells within these cell states tend to behave differently in terms of cell-cell communication, thus leading to a greater information loss when averaging CCC over the cell type (Fig. **2I**). Similarly, several CCC pathways such as non-canonical WNT and EPHB have high heterogeneity scores, implying that cells with different CCC behavior are spread heterogeneously across different cell states (Fig. **2J**). In general, the higher the heterogeneity score, the higher the loss in information when averaging CCC over cell types, thus motivating the modeling of CCC at the meta-cell resolution.

### Experimental validation by analysis of downstream regulations of the EGF pathway in skin-equivalent organoids

Next, we use the FibHSE keratinocytes^33^ to validate scRICH’s predictions about CCC heterogeneity and CCC pathway co-dependence experimentally. These organoids reproduce the stratification of basal keratinocytes (BAS) that differentiate into spinous (SPN), granular (GRN), and cornified (CRN) keratinocytes found in the epidermis (Fig. **3A**). We focus on the EGF CCC pathway that was previously implicated in keratinocyte differentiation^33^ and exhibits a Sender, Sender/Receiver, and Weak signaling modes (see Fig. **2B-C**). We identify multiple cell states including SPN-1 and SPN-3 that exhibit high EGF heterogeneity scores. Therefore, these cell states are mixtures of EGF Sender, Sender/Receiver, and Weak cells (Fig. **3B-C**).

**Figure 3.**
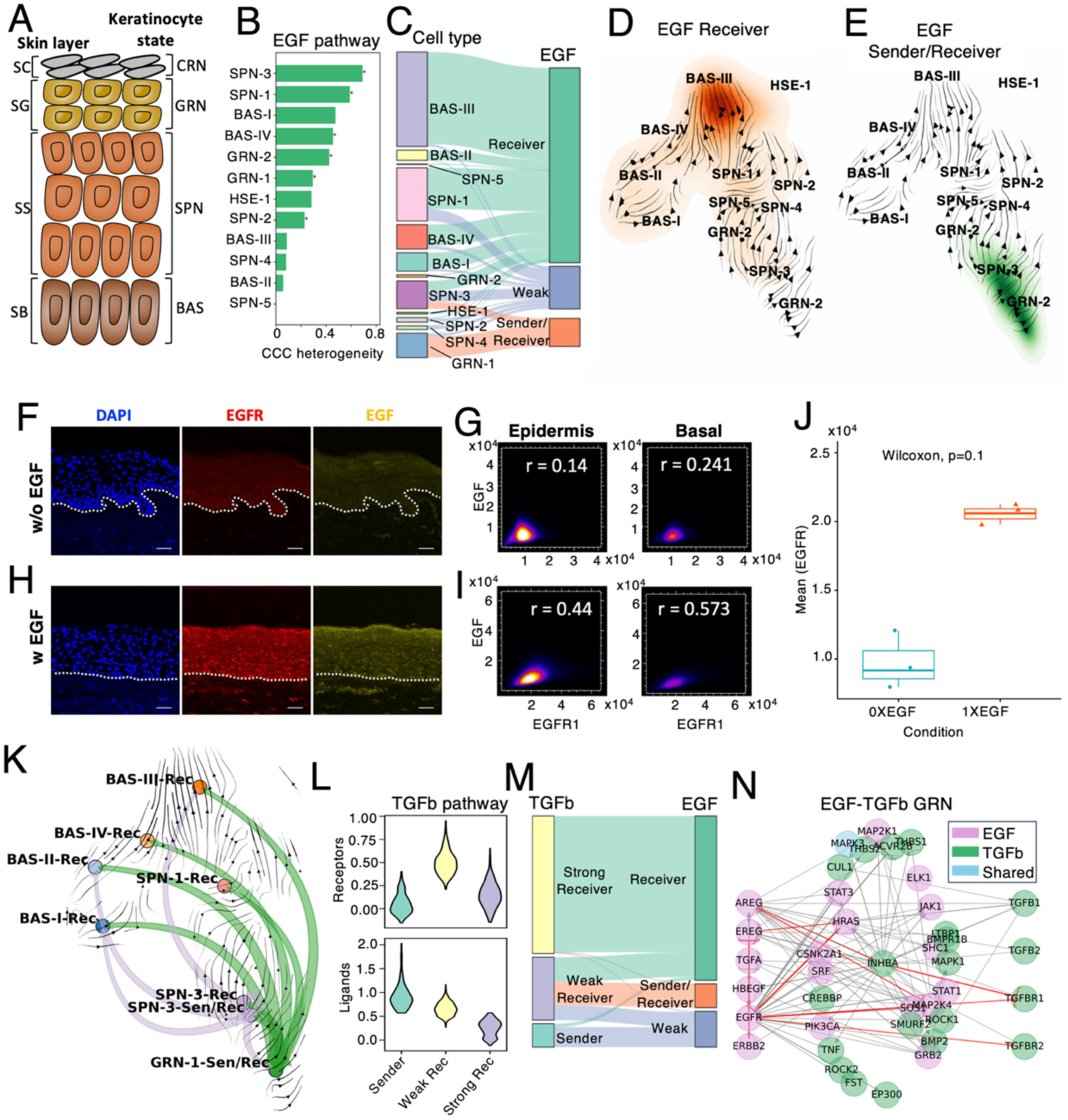
The EGF and TGFB cell-cell communication in FibHSEs. **(A)** Epidermal differentiation in human skin. Keratinocytes transition from basal (BAS; early) to spinous (SPN) and granular (GRN; late) states as they migrate from the stratum basale (SB) through stratum spinosum (SS) and stratum granulosum (SG), ultimately reaching the cornified (CRN; terminal) stage in the stratum corneum (SC). **(B)** The CCC heterogeneity of the EGF pathway in the keratinocyte cell states. Asterisk indicates statistical significance of permutation testing. **(C)** Mapping between keratinocyte states and EGF CCC signaling modes. **(D-E)** Density of EGF Receiver and EGF Sender/Receiver cells in low dimensional UMAP space superimposed with RNA velocity **(F)** Immunofluorescence of the epidermis layer in FibHSE showing cell nuclei, EGF, and EGFR layer without EGF treatment. **(G)** Correlation between EGF and EGFR in the epidermis and basal layer without EGF treatment. r indicates Pearson’s correlation coefficient. **(H-I)** Same as (F-G) with EGF treatment. Scale bars, 50μm. Dashed lines denote the epidermal-dermal junction. **(J)** Quantification of EGFR expression in the epidermis based on replicates and EGF dosage. **(K)** The EGF CCC network between keratinocyte subsets and its relation to cell state transitions. **(L)** The TGFB signaling modes in the FibHSE keratinocytes (gene expression normalized and log-transformed). **(M)** Mapping between TGFB and EGF CCC modes. **(N)** The gene regulatory network connecting the EGF and TGFB pathways, whereby red edges indicate the path of max flow connecting EGF and TGFb receptors.

In particular, cells in the EGF Sender/Receiver mode are found across several different keratinocyte cell states, and are especially represented in the terminal states including SPN-3 and GRN-1. Conversely, the EGF Receiver mode is mainly observed at earlier stages of the keratinocyte lineage, and especially in basal states including BAS-III, BAS-IV, and SPN-1 (Fig. **3D-E**). To confirm the predicted role of EGF Sender/Receiver cells, we performed immunostaining for the EGF ligand and the EGFR receptor in FibHSE organoids. EGF and EGFR were lowly expressed without EGF induction, but became much highly expressed with soluble EGF in solution (Fig. **3F, H**). Furthermore, exposure to external EGF ligand increased the co-localization of the EGF and EGFR in the organoids, showing that their expression correlates both in the basal layer as well as whole epidermis (Fig. **3G, I, J**). Overall, these observations support a model whereby late-stage keratinocyte relay information via EGF to early-stage keratinocyte, which is specifically visualized in scRICH with a superposition plot that overlays the cell state transition field and EGF-CCC (Fig. **3K**).

Next, we apply scRICH to model computationally the connection between EGF and other CCC pathways involved in EMT regulation including TGFB, NOTCH and CDH. First, we identify three TGFB CCC signaling modes including a Sender, strong Receiver and weak Receiver (Fig. **3L**), and compare cell behavior in the two pathways. Interestingly, Strong TGFB receivers tend to be EGF Receivers. TGFB Senders, however, are mostly weak with respect to EGF. Finally, the TGFB weak receptor mode is almost equally spread among the three EGF signaling modes (Fig. **3M**). We further investigate the strong correlation between several EGF and TGFB signaling modes by inferring the intracellular gene regulatory network. Specifically, we reconstruct the network connecting the EGF ligands/receptors including EGFR, AREG, EREG, TGFA, HBEGF, and ERBB2 with the TGFB ligands/receptors including TGFB1, TGFB2, TGFBR1, and TGFBR2. By applying a maximum flow algorithm to the predicted GRN, scRICH predicts the top connections and intermediate mediators connecting the EGF and TGFB CCC signaling modes, which include HRAS and INHBA (Fig. **3N**), both of which were previously linked to EMT^34–36^. Consistently, the expression of HRAS and INHBA increased upon exposure to exogenous EGF in our experiment (Supplementary Fig. **S3**), suggesting a connection between the EGF sender/receiver communication mode, HRAS/INHBA activation, and TGFB signaling. To better evaluate the relation between EGF signaling and EMT during keratinocyte differentiation, we evaluate CDH (Cadherin) signaling that directly regulates cell adhesion, identifying three CCC modes corresponding to weak, medium and string adhesion (Supplementary Fig. **S4A**). Interestingly, EGF communication modes were strongly correlated with cell adhesion: while EGF weak cells are mostly CDH weak as well, EGF Sender/Receivers mostly exhibited strong adhesion (Supplementary Fig. **S4B**), and EGF signaling was predicted to directly induce cell adhesion via CDH4 activation in the inferred EGF-CDH regulatory network (Supplementary Fig. **S4C**). This prediction resonates well with the co-localization of EGF and CDH4 observed experimentally via immunostaining, whereby CDH4 expression is upregulated upon EGF induction (Supplementary Fig. **S4HD-F**). A similar analysis of the Notch CCC pathway highlights once again very strong correspondence between EGF and NOTCH CCC signaling modes. Specifically, EGF Sender/Receivers are almost exclusively NOTCH Receivers, whereas NOTCH Sender/Receivers are almost exclusively EGF Receivers. Finally, low activity for the two pathways goes hand in hand (Supplementary Fig. **S4G-H**). The emerging correlation between the EGF and NOTCH receiver modes was supported by the multiple activation paths identified in the predicted EGF-NOTCH gene regulatory network (Supplementary Fig. **S4I**), including the intermediate regulator STAT1 that increases along with both NOTCH and EGF (Supplementary Fig. **S4J-M**).

Overall, these results highlights that the CCC behavior in FibHSEs is quite heterogenous, leading to loss of information when evaluating at the level of cell types. Conversely, scRICH identifies well defined signaling modes of cells with coherent communication behavior and clear relationships between distinct CCC pathways.

### ScRICH distinguishes subsets of sender and receiver cells within cell types

To demonstrate scRICH’s ability to identify within-type CCC heterogeneity, we analyze a scRNA-seq skin wound healing dataset that includes multiple distinct types of fibroblasts, keratinocytes, and immune cells (Fig. **4A**). An overview based on either ligand/receptor average expression or number of detected ligands/receptors in the dataset suggests the activation of several CCC signaling pathways (Fig. **4B-C**).

**Figure 4.**
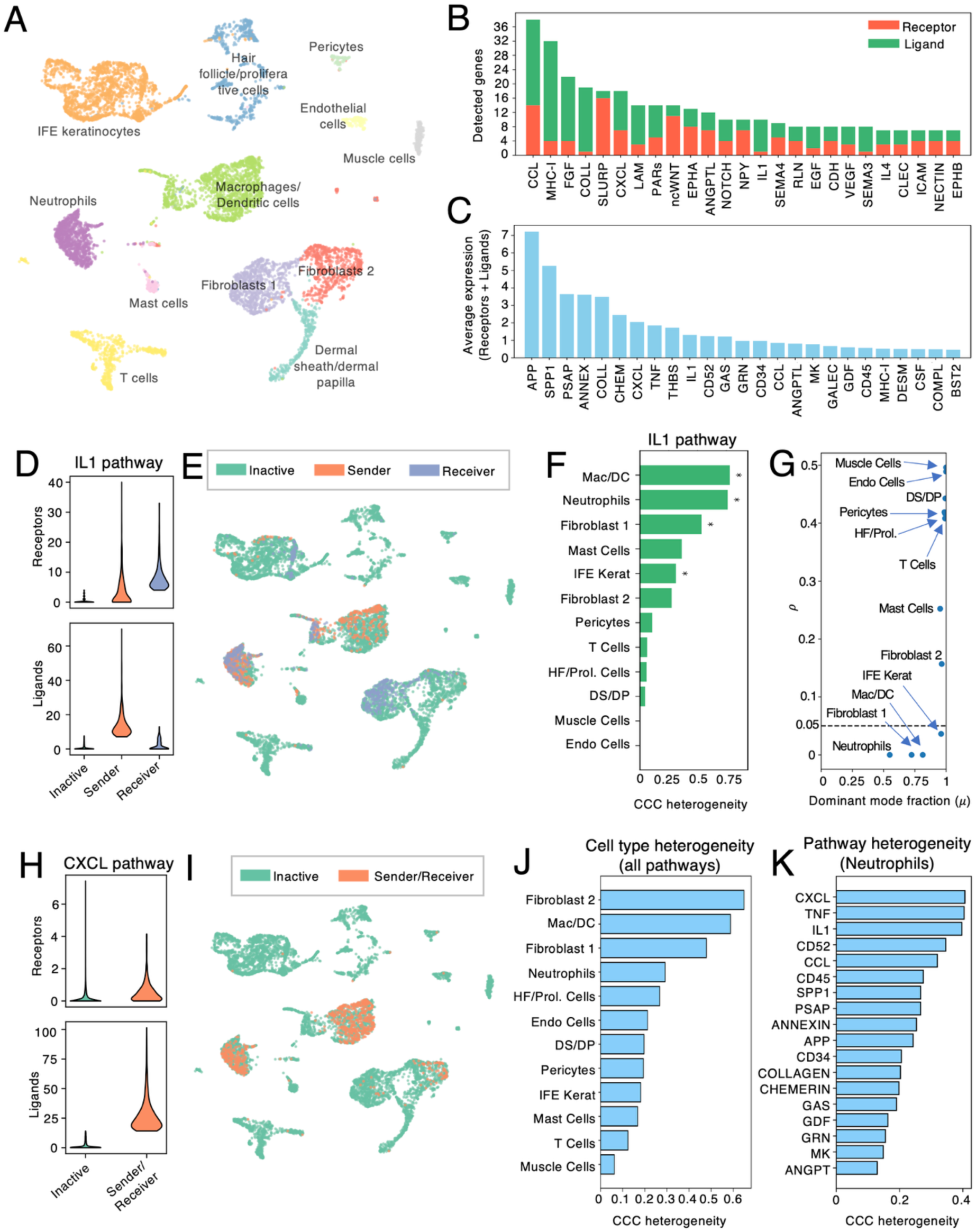
Functional separation of cell types into robust CCC modes in skin wound healing. **(A)** UMAP representation of the wound healing cell types. **(B-C)** The top CCC signaling pathways identified skin wound healing based on number of detected genes (B) and average expression of ligands and receptors (C). **(D)** The IL1 communication modes corresponding to an inactive (green), sender (orange), and sender (blue) states. **(E)** Low-dimensional embedding whereby cells are color-coded with IL1 CCC communication modes. **(F)** The CCC heterogeneity of the IL1 pathway. Asterisk indicates statistical significance of permutation testing. **(G)** Permutation testing p-value as a function of fraction of cells in the dominant CCC signaling mode in each cell type. Cell types below the threshold of 0.05 are selected for functional separation. **(H-I)** Same as (D-E) for the CXCL pathway. **(J)** Cell type CCC heterogeneity score (average over all CCC pathways). **(K)** Heterogeneity score of each CCC pathway in the Neutrophil cell type.

To highlight a representative workflow, we focus on the IL1 cell-cell communication pathway. IL1 pathway is associated to inflammation response and thus plays an important role in coordinating cell behavior during wound healing^37^. When clustering cells based on IL1 ligand/receptor expression, we identify 3 clusters associated to a Receiver mode (high receptor, low ligand expression), a Sender mode (low receptor, high ligand expression), and Inactive mode (low expression of both) (Fig. **4D**). Given scRNA-seq data sparsity, it could be hypothesized that the Inactive mode originates from cells with low RNA counts. Visualizing cell membership to the 3 IL1 modes in the UMAP embedding, however, highlights a specific pattern whereby 4 cell types (Fibroblast 1, Neutrophils, Macrophage/Dendritic Cells, and IFE Keratinocytes) possess subgroups of Sender and Receiver cells, whereas all other cell types are almost exclusively Inactive (Fig. **4E**). Therefore, these cell types exhibit high IL1 heterogeneity scores (Fig. **4F**). To proceed with functional separation of these cell types between signaling modes, we perform permutation testing (**Fig. 4G**), indicating that these 4 cell types should be split for functional characterization of IL1 cell-cell communication. Specifically, the Fibroblast1 and keratinocyte cell types are composed by large populations of IL1-inactive cells and smaller but well-separated populations of IL1-Receiver cells. Therefore, averaging IL1-based communication inference over cell types would weaken the IL1-receiver behavior. Most interesting, the Neutrophil and Macrophage/Dendritic cell types include well-separated subsets of IL1-Receives and IL1-Senders. Therefore, cell type-level averaging would miss this separation, instead predicting a Sender/Receiver behavior. Permutation testing allows to select robust patterns in CCC heterogeneity. For example, Mast cells exhibit a high IL1 heterogeneity score (see Fig. 4F). The low Mast cell count, however, prevents a statistically robust functional separation within this cell type. In other words, the null hypothesis that a small subset of IL1-Sender cells exhibits a IL1-Sender, rather than a IL1-Inactive mode by random chance cannot be rejected with statistical significance. To showcase the importance of intra-type heterogeneity, we consider the CXCL pathway that is exclusively activated in Neutrophil and Macrophage/Dendritic cells. Compared to IL1, CXCL features a true Sender/Receiver state whereby individual cells co-express CXCL receptors and ligands (Fig. **4H-I**).

After identifying CCC signaling modes for all the most expressed CCC pathways, each cell type can be characterized by a CCC heterogeneity score that summarizes the coherence of CCC behavior of cells of same type across many CCC pathways (Fig. **4J**). Fibroblasts (both type-1 and 2), Macrophages/DC cells, and Neutrophils exhibit more heterogeneous behavior (high CCC heterogeneity score), whereas muscle cells, T-cells, and Mast cells exhibit more coherent CCC behavior (low CCC heterogeneity score). Individual cell types can be further characterized based on their CCC heterogeneity score for each distinct pathway. For example, Neutrophils behave heterogeneously along specific pathways including CXCL, TNF, IL1, and CD52 (Fig. **4K**).

### ScRICH identifies cell-cell communication gradients along cell differentiation lineages

To further test ScRICH’s ability to capture CCC evolution during cell fate transitions, we investigate cell differentiation during erythroid cell maturation where blood progenitor cells differentiate into erythroid cells^38^ (Supplementary Fig. **S5A**). As blood progenitor cells undergo enucleation, we first inspected the presence of unspliced RNA, confirming active transcription in all cell types (Fig. **S5B**). We identify several CCC pathways that are highly expressed and showcase heterogeneous behavior across erythroid cell states including IGF and VEGF (Fig. **5A**, Supplementary Fig. **S5C-E**). Specifically, we identify three IGF CCC modes including a sender, receiver, and inactive (Supplementary Fig. **S5F-H**).

**Figure 5.**
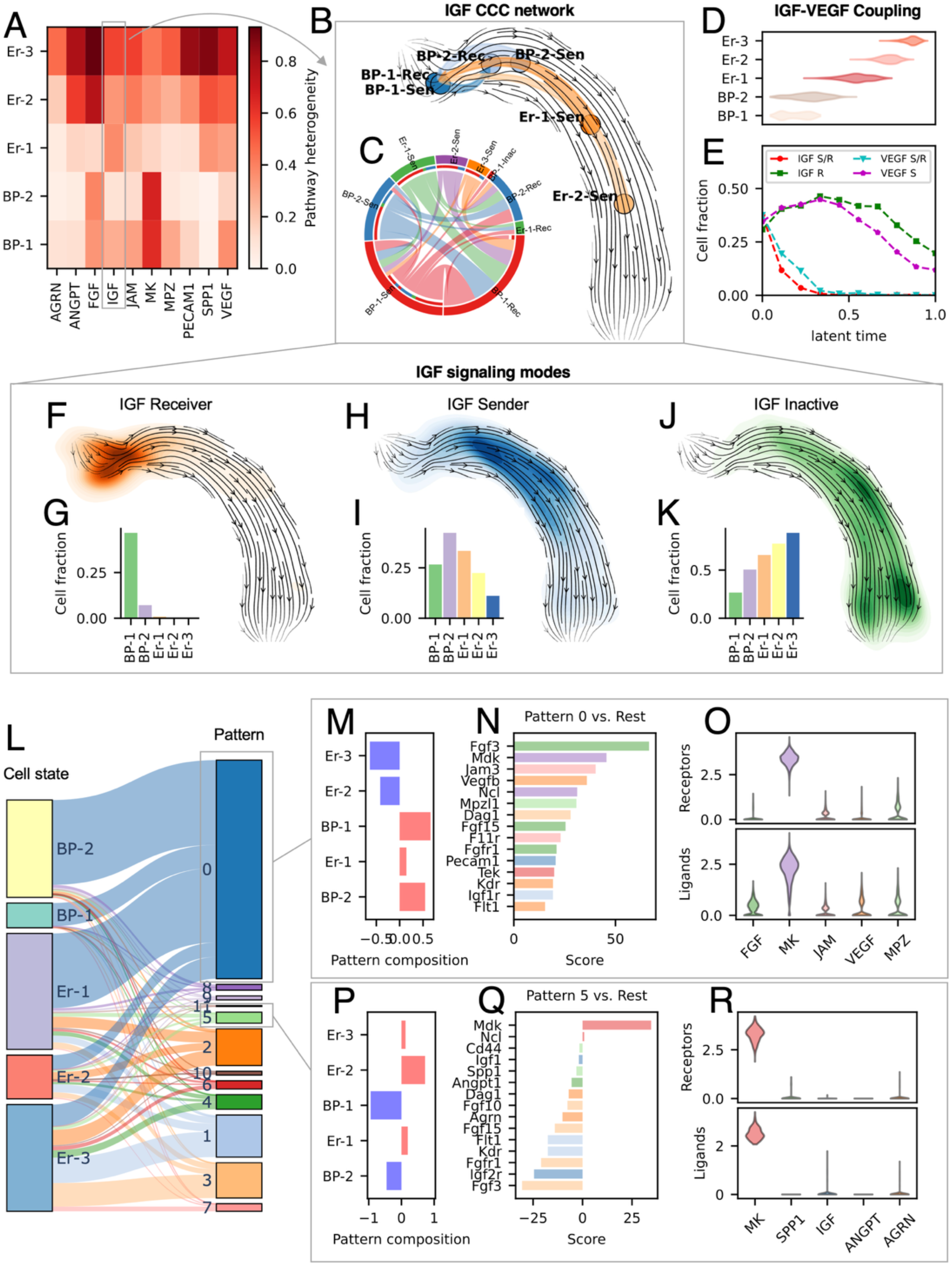
Application of ScRICH to the erythroid lineage. **(A)** The CCC heterogeneity of erythroid cell states for multiple CCC pathways. **(B-C)** The IGF CCC network depicted over RNA velocity (B) and as chord diagram (C). **(D)** The distribution of cell state over latent time. **(E)** The fraction of IGF and VEGF CCC modes as a function of latent time. **(F)** The cell density plot of IGF receivers. **(G)** Fraction of IGF receivers in the erythroid cell states. **(H-I)** Same as F-G for IGF senders. **(J-K)** Same as F-G for IGF inactive. **(L)** Sankey diagram of matching between erythroid states and CCC patterns. **(M)** The representation fold-change of erythroid cell states in CCC pattern 0. **(N)** The top 15 ligands and receptors expressed in CCC pattern 0. **(O)** Violin plot of the top 5 receptors (top) and ligands (bottom) in CCC pattern 0. **(P-R)** Same as M-O for CCC pattern 5.

Juxtaposing the IGF communication modes with the directions of cell state transition computed with UnitVelo^39^ suggests a strong interaction between the IGF CCC network and cell fate. Here, Unitvelo is chosen because it was previously shown to generate more robust cell transition trajectories compared to scVelo. scRICH, however, is compatible with both packages. The early-stage BP-1 cells split into subpopulations of BP-1 senders and BP-1 receivers, whereas the later Er-1 and Er-2 cells are mostly IGF senders, overall suggesting a transition mechanism whereby later stage erythroid cells communicate with early-stage cells and may contribute to their differentiation (Fig. **5B-C**). The evolution of IGF along the lineage is further captured by inspecting the fraction of cells in different IGF CCC modes along latent time, which shows a progressive decrease of cells that behave as receivers of both IGF and VEGF (Fig. **5D-E**). This trend is further visualized by cell density plots whereby early cells tend to behave as IGF receivers, whereas cells in the intermediate lineage mostly behave as senders, and finally IGF signaling is mostly inactive in the later stages (Fig. **5F-K**).

Finally, we investigate the emerging trend in CCC using unsupervised learning strategy, leading to the identification of 12 CCC patterns with different levels of representation in the erythroid cell states (Fig. **5L**). These patterns exemplify how different CCC pathways are orchestrated at different stages of the erythroid lineage. For example, pattern 0 represents a CCC behavior over-represented in early-stage cells including BP-1, BP-2, and Er-1, featuring strong expression of the Midkine (MK) receptor – a growth factor related to several cellular programs^40^ – and the FGF, MK, and VEGF ligands among others (Fig. **5M-O**). In other words, albeit belonging to different cell states, cells in the pattern 0 share a common CCC behavior when evaluating their communication profile across several different CCC pathway. Conversely, pattern #5 is enriched of cells in the later Er-2 stage and features co-expression of the MK ligand and receptor (Fig. **5P-R**). Overall, these CCC patterns provide a comprehensive representation of how different CCC pathways are orchestrated together at different stages of the cell lineage.

### TGFB and BMP modulate parallel cell state transition trajectories

To test ScRICH across different biological scenarios, we next consider a time course of EMT in the cancer cell line OVCA420 under exogenous TGF*β* exposure^41^. Cells in the dataset undergo EMT from 0 days to 7 days of TGF*β* exposure as reported based on cell morphology in the original publication as well as by examination of signatures of epithelial and mesenchymal gene sets (Supplementary Fig. **S6A-B**). Clustering cells based on gene expression rather than time point suggested 3 cell states corresponding to epithelial, intermediate and mesenchymal cell states, with the mesenchymal state becoming increasingly populated at later time points, in good agreement with the reported EMT induction (Supplementary Fig **S6C-E**). The TGF*β* pathway, a well-known driver of EMT, splits into 3 CCC signaling modes including TGF*β*-receiver, TGF*β*-sender, and TGF*β*-inactive (Fig. **6A**). While the TGF*β* CCC modes were spread heterogeneously across the E, I, and M cell states, the fraction of TGF*β* sender cells increased along the pseudotime axis, suggesting that cells undergoing EMT could in turn induce EMT in neighboring cells via TGF*β* signaling (Supplementary Fig. **S6F-H**). By overlaying the TGF*β* signaling states and cell state transition trajectories marked by RNA velocity, ScRICH suggested the cells in the early stage of the EMT trajectory tend to be mostly (epithelial, TGF*β*-inactive) or (epithelial, TGF*β*-receiver); conversely, at later stages cells switch to the (intermediate, TGF*β*-receiver) and finally to the (mesenchymal, TGF*β*-sender) states (Fig. **6B-D**).

**Figure 6.**
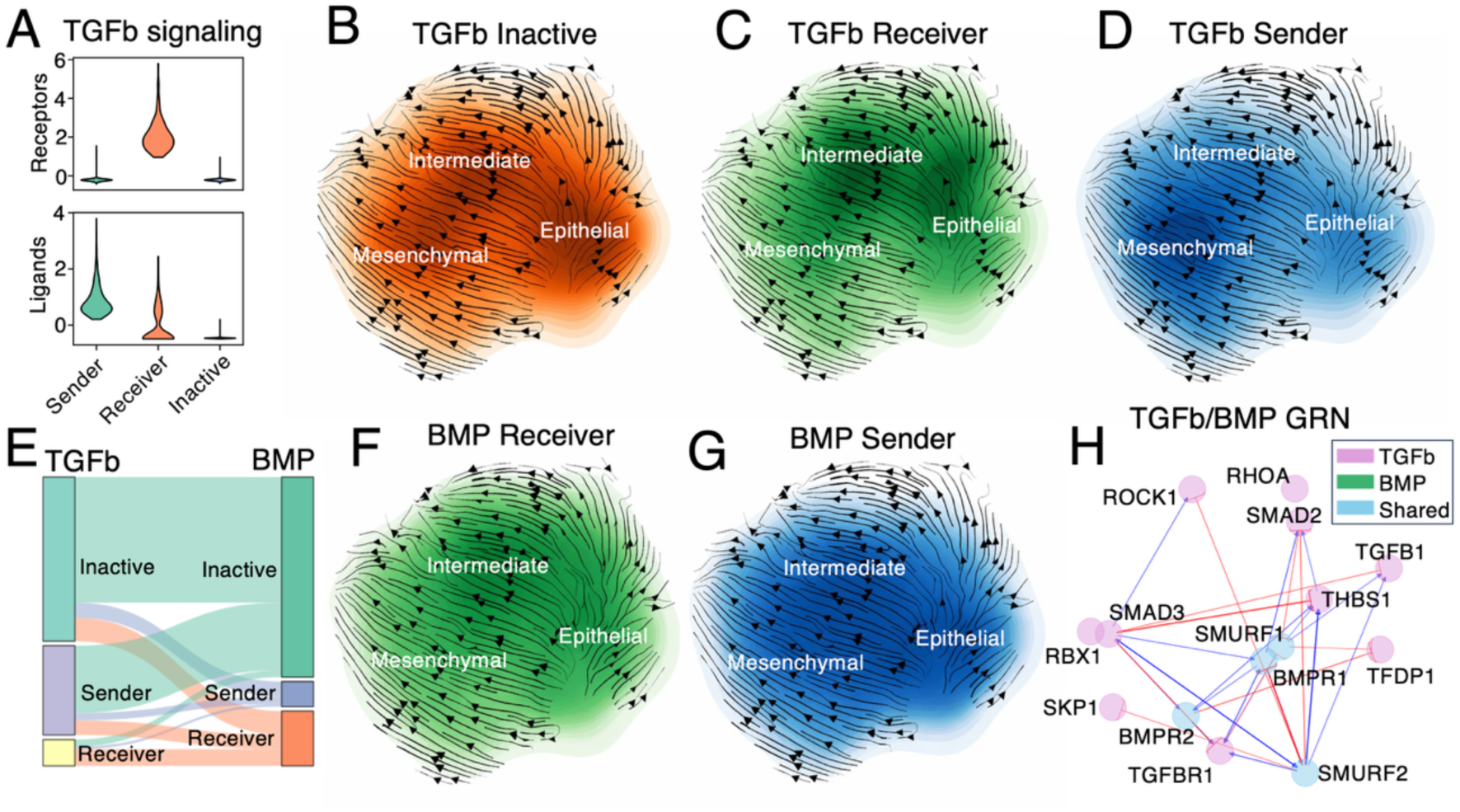
Interplay of TGFb and BMP during epithelial-mesenchymal transition. **(A)** The TGFb signaling modes in OVCA420 cells (expression is normalized and log-transformed). **(B-C-D)** The cell density of TGFb CCC modes overlayed on RNA velocity for TGFb inactive (B), receivers (C), and senders (D). **(E)** Sankey diagram of correspondence between TGFB and BMP CCC modes. **(F-G)** Cell density of BMP receivers and BMP senders. **(H)** The joint TGFb-BMP gene regulatory network. Purple and blue shading represent nodes associated to the TGFb pathway or shared between the two pathways, respectively.

RNA velocity suggests that epithelial cells can either transition directly to the mesenchymal state or traverse a “longer” path through the intermediate state. To characterize the signaling mechanism along these trajectories, we evaluated the pathway similarity, finding a high mutual information between TGF*β* and BMP, another pathway usually associated with EMT (Supplementary Fig. **S6I**). Similar to TGF*β*, BMP exhibited both a sender and receiver signaling mode in the dataset (Supplementary Fig. **S6J-K**), which were highly correlated with the TGF*β* signaling modes. Specifically, cells that were TGF*β*-inactive and TGF*β*-receiver had high probability to be BMP-inactive and BMP-receiver, respectively (Fig. **6E**). Interestingly, TGF*β* and BMP showed different activation patterns along the EMT trajectory. The (mesenchymal, TGF*β*-sender) cells tend to be largely BMP-inactive, whereas the (intermediate, TGF*β*-receiver) cells tend to be BMP-receiver as well. Combined with the RNA velocity transition trajectories, these results suggest alternative EMT trajectories. First, a direct E-M trajectory that relies on TGF*β* but not BMP activation; second, a E-I-M trajectory where both TGF*β* and BMP are activated (Fig **6F-G**). To characterize the relationship between TGF*β* and BMP further, we reconstructed the intracellular GRN connecting the TGF*β* and BMP pathways, which identified several transcription factors commonly associated with EMT including SMAD3, SMAD6, and SMURF2 (Fig. **6H** and Supplementary Fig. **S6M-N**).

### ScRICH identifies spatial patterns of sender and receiver cells from spatial transcriptomics data

Finally, we consider a spatial transcriptomics dataset of chicken heart development^42^ originally obtained via the 10x Genomics Spatial RNAseq Visium platform to leverage the spatial organization of cell-cell communication. While the spatial distribution of cell types in this dataset is highly heterogeneous (Supplementary Fig. **S7A**), we hypothesize that CCC should still exhibit spatial organization due to limit in the diffusion of ligands across the tissue. Among several CCC pathways (Supplementary Fig. **S7B**), we focus specifically on the highly expressed MK pathway, which is heavily implicated in the regulation of stem cell renewal, angiogenesis, and inflammation^43^. Strikingly, we observe a robust spatial organization when classifying cells based on their MK communication behavior instead of cell type. Specifically, the three identified MK communication modes including receiver, sender/receiver and weak sender/receiver are distributed heterogeneously across cell types (Fig. **7A**) but are mainly localized in distinct spatial locations (Fig. **7B-D**), resulting in a CCC network whereby multiple cell types contribute to the signaling (Fig. **7E**). To further summarize the spatial organization of the MK CCC pathway, we define a spatial heterogeneity index which quantifies whether cells assume a CCC communication mode similar to their spatial neighbors. Interestingly, cells in different spatial locations exhibit varying degrees of spatial heterogeneity, with the region corresponding to the strong sender/receiver MK mode being the more robust/less heterogeneous, whereas regions corresponding to weakly interacting cells are more heterogeneous (Fig. **7F**).

**Fig. 7.**
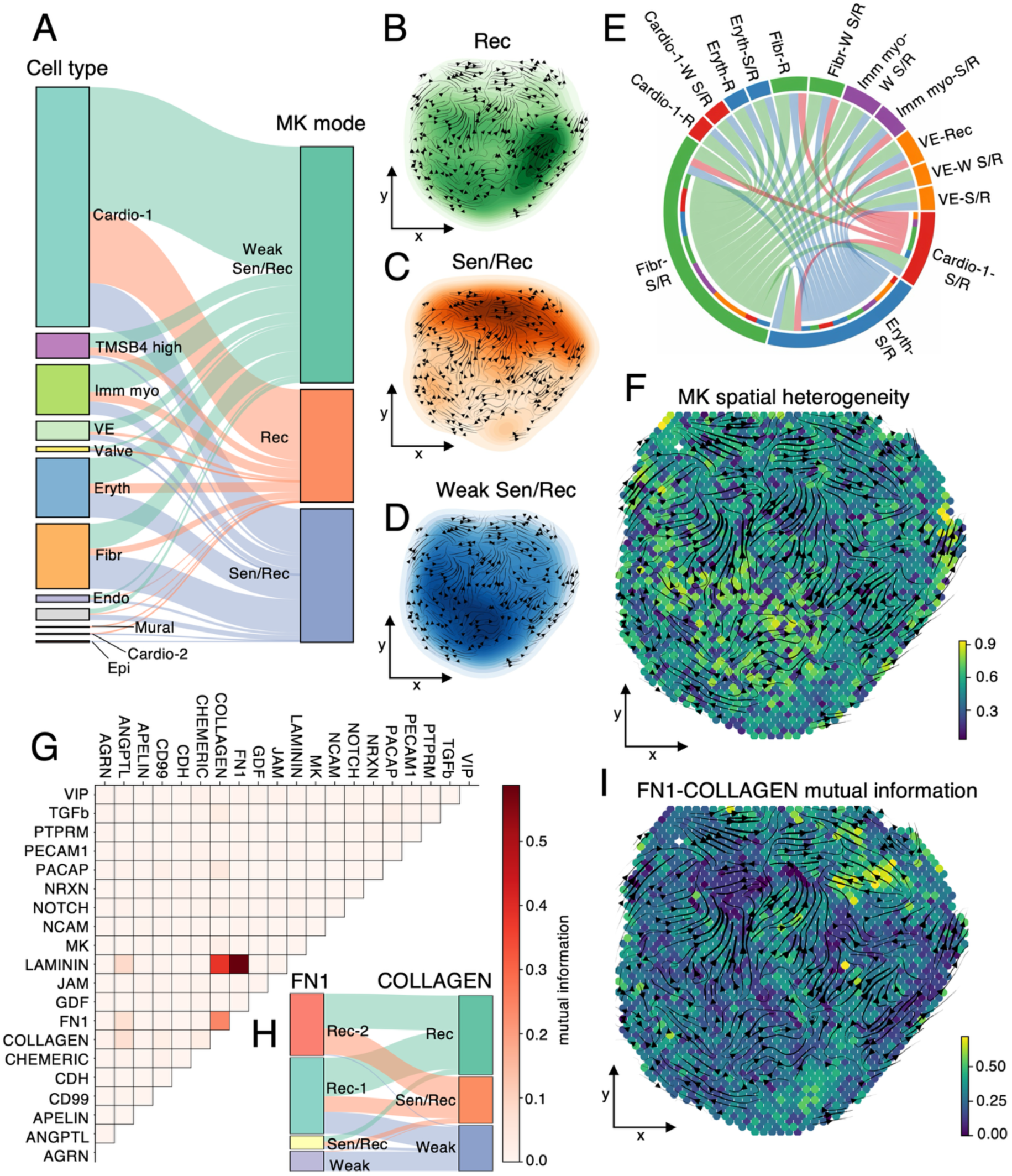
The spatial organization of cell-cell communication modes in the developing chicken heart. **(A)** Composition of cell types and MK CCC modes. **(B-C-D)** Spatial distribution of the identified MK signaling modes including MK receiver (B), MK sender/receiver (C), and MK weak sender/receiver (D). **(E)** The MK communication network. **(F)** The spatial heterogeneity of MK CCC signaling. **(G)** Mutual information between CCC pathway. **(H)** Correspondence of CCC signaling modes between FN1 and Collagen pathways. **(I)** Spatial distribution of the FN1-Collagen mutual information in the developing chicken heart.

The spatial segregation of cells with distinct MK communication behaviors suggests that cell-cell interactions play a significant role in the development of the tissue. Thus, we investigate whether this spatial organization is also observed in the correlations between CCC pathways. Based on mutual information analysis, we identify pairs of pathways that are highly correlated, such as FN1 and COLLAGEN (Fig. **7G-H**). To test the relationship between these CCC pathways in different anatomical regions of the tissue, we define a local, space-dependent mutual information that only considers the relation between the pathways in the neighborhood of a specific spatial location. This analysis showcases different anatomical locations of the tissue where the relationship between the CCC pathways differs (Fig. **7I**). For example, certain regions are characterized by high mutual information, thus implying that it is possible to predict a cell’s communication behavior in the COLLAGEN pathway if the FN1 behavior is known, or *vice versa*. Generally, the connection between these pathways is not surprising as both FN1 and COLLAGEN play a key role in the processing of interaction between the cell and the extracellular matrix^44,45^. These pathways, however, have low mutual information in specific spatial regions, potentially suggesting distinct roles locally in tissue development.

## Discussion

Cell-cell communication (CCC) enables the flow of information in multicellular systems and connects the tissue and cellular scales through intricated gene regulatory networks (GRN) that process external information and inform cellular responses, for example by activating secondary CCC pathways. High throughput single cell and spatial transcriptomics technologies offer the unprecedented opportunity to observe these biological processes at single cell resolution^15,46,47^. Yet, existing models mostly focus on the scales of CCC mediated by ligand-receptor binding and intracellular GRN separately^1,48,49^, thus failing to provide a holistic picture of how CCC and cellular identity interact and evolve together in complex cell lineages.

To address this existing gap, we have developed *scRICH*, a mathematical framework and bioinformatics package that leverages single cell transcriptomics, prior biological information on CCC pathways, and mathematical modeling of inter- and intracellular signaling dynamics. *scRICH* takes advantage of the unspliced and spliced RNA counts from single cell RNA sequencing (scRNA-seq) data to (1) Quantify the diverse CCC behaviors and heterogeneity within cell types, (2) Build a robust model of cell-cell communication that couples ligand-receptor binding with intracellular GRN response, thus enabling to investigate the association between distinct CCC pathways, and (3) Identify emerging global communication patterns and their evolution during cell fate transition events.

Recently, FlowSig proposed a learning framework to discover relations between signaling pathway activation and expression of new ligands/receptors using conditional independence^50^. Compared to this method, scRICH focuses on (1) the heterogeneity of CCC behaviors within cell types, and (2) the relation between CCC and cell fate by incorporating RNA splicing dynamics, thus providing unique and complementary insight. First, scRICH treats cell types and CCC behavior as independent observations, thus enabling to quantify how cells of same type operate via different (sets of) CCC pathways, and building CCC networks at the intermediate scale of meta-cells, i.e., groups of cells that share the same type and communication behavior. This approach enabled us to identify cell sub-populations that operate via distinct sets of cell-cell interactions despite sharing the same set of core marker genes, such as groups of EGF sender/receiver spinous cells that was validated experimentally based on EGF ligand-receptor co-expression in FibHSE organoids. Second, scRICH employs multilayer networks and mutual information to predict the connection between CCC pathways, for example predicting intermediate transcription factors HRAS and INHBA that orchestrate a joint (EGF receiver, TGFB receiver) cellular state in basal keratinocytes. When coupled with cell fate transition information, this approach can clarify how CCC pathways work together to modulate early decisions in cell fate. For example, scRICH predicts parallel EMT trajectories driven by TGFB in ovarian cancer-derived cells, whereby a first trajectory shows strong coupling between the TGFB and BMP pathways and passes through an intermediate epithelial/mesenchymal cell state, whereas a second, BMP-independent trajectory directly connects the E and M cell states.

Despite these encouraging results, this method could be generalized in the future to interface other Omics data modalities besides scRNA-seq to improve predicting power. First, we follow a common assumption in the field and utilize RNA counts to estimate ligand/receptor expression, thus disregarding several biological steps including translation and ligand/receptor transport to cell surface. This approximation poses specific challenges in biological contexts or datasets with poor protein-RNA correlation. Conversely, proteomics^51^ or CITE-seq – which measures protein expression at cell surface^52,53^ – could provide better estimation. Similarly, GRN inference based exclusively on single cell transcriptomics, albeit by integrating unspliced and spliced RNA layers, would benefit in the future from the integration with DNA accessibility data. For example, scATAC-seq has been used in combination with regular (i.e., spliced) scRNA-seq for robust GRN inference by providing prior on whether DNA sequences associated to specific genes are accessible^54,55^. Moreover, scRICH leverages gene expression and/or RNA velocity (when available) of downstream targets to control for false positive, thus detecting CCC if and only if a downstream response is observed in the receiver cell. More strategies, however, should be investigated in the future to further alleviate false positives. For example, scRICH currently employs spatial information (ST) exclusively to investigate the spatial organization of CCC modes within a tissue, whereas other methods employ ST to cut-off interaction between faraway cells^56–60^, and the inferred results could be applied to non-spatial scRNA-seq datasets with more cells and deeper sequencing. For example, stMLnet combines ST for false positive correction and intracellular gene signature for CCC prediction^61^. Finally, an additional limitation of ST is the increased data sparsity, as sequencing-based, whole-genome data is needed as scRICH considers ligands, receptors, and downstream targets over several CCC pathways. Future applications of scRICH will benefit from integration of ST with non-spatial data for improved data quality^62^.

In addition to multi-modal and/or spatial high-throughput data, scRICH could be integrated with dynamical modeling of cell fate^63–66^ and/or temporal information including time course and lineage tracing to further explore the relation between cell fate and cell-cell communication. Existing methods leverage these types of data to estimate cell state transitions under the assumption of cell autonomous processes. For example, Cospar integrates lineage tracing and single cell transcriptomics to build a Markov chain Monte Carlo model of cell-cell and state-state transitions^67^, while TIGON implements optimal transport to estimate transitions and cell growth between consecutive points in time course data^68^. Developing non-autonomous models that integrate multiscale CCC in the style of scRICH with cell transition states would provide a unifying framework to understand the influence of cell-cell interactions on cell-fate decisions.

## Materials and Methods

1. **Input data and preprocessing.** The input to scRICH is the scRNA-seq data object, which is assumed to be preprocessed following standard preprocessing pipelines^69,70^, and includes log-normalized RNA counts, filtered genes and cell annotations. In addition, the unspliced and spliced RNA layers are required for the downstream modeling, GRN inference, and visualization. The unspliced/spliced counts are obtainable via compatible python packages including velocyto^31^ and kallisto bustool^71^ to allow efficient streamline of the analysis in a single notebook.

2. **Cell-Cell signaling database.** scRICH includes a pre-built database comprising a list of ligands and receptors for cell-cell communication pathways updated based on CellChat database^9^, spanning both human and mouse. Moreover, a database of downstream transcription factors (TF) was constructed by merging information about ligand-receptor-TF connections from NicheNet^16^ and exFinder^25^. All scRICH’s analysis steps can be executed from the pre-built database. Additionally, several package applications can be integrated with custom-built lists of upregulated/downregulated downstream genes for individual CCC pathways.

3. **Average expression within single cells.** Several modeling steps of scRICH use average expression of ligands, receptors, and/or downstream targets within individual cells. scRICH offers multiple possibilities to compute these averages, which can be chosen by the user, including: (1) Log-normalized RNA counts (standard output of scanpy’s preprocessing^69^); (2) moments of RNA counts; and (3) Average over the cell neighborhood graph.

4. **Identification of cell-cell signaling modes.** For each pathway included in the database, scRICH identifies the number of communication modes by applying symmetric non-negative factorization of a cell-cell similarity matrix^72^, which has been previously applied for unbiased clustering of scRNA-seq data^29,73^. First, different K-means cell clustering solutions are computed based solely on the average expression of ligands and receptors in each cell. By default, n=5 clustering solutions with randomized seed are computed for k=3,4,5,6, whereby all parameters can be modified by the user. Second, a cell-cell similarity matrix is computed where a matrix element *S*_*ij*_ ∈ [0,1] is the fraction of clustering solutions where cells *i* and *j* are grouped in the same cluster. The similarity matrix is then decomposed using symmetric non-negative factorization:

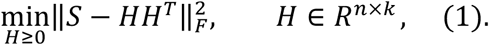

Where *S* is the cell-cell similarity matrix, *H* is a non-negative low-rank matrix, and *n*, *k* are the number of cells and clusters. The optimal number of signaling modes is identified based on the largest gap between the sorted eigenvalues of the matrix H.

5. **Quantification of cell-cell signaling heterogeneity.** After identification of CCC modes of a given pathway *P*, the dataset can be categorized into *N* cell types and *K* CCC modes, resulting in *N* × *K* sub-clusters. The heterogeneity (ℎ) of cell type (*i*) with respect to the CCC pathway *P* is defined as:

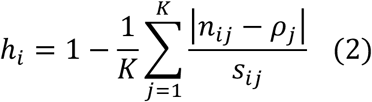

Where *n*_*ij*_is the fraction of cells in cell type *i* that were clustered in CCC mode *j*, and 𝜌_j_ is the overall fraction of cells in signaling mode *j* in the entire dataset. Finally, *s*_*ij*_is a normalization factor to ensure that ℎ_*i*_is bound between 0 and 1:

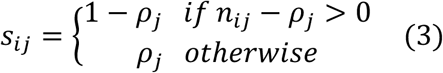

Based on this definition, the cell type heterogeneity score ℎ_*i*_ is minimized (ℎ_*i*_ = 0) when all cells within cell type *i* assume the same CCC mode; conversely, ℎ_*i*_ is maximized (ℎ_*i*_ = 1) when cells within cell type *i* are divided between all CCC modes with same proportions observed in the whole dataset.

An overall heterogeneity score for the CCC pathway *P* is obtained by averaging the heterogeneity scores ℎ_*i*_ over all cell types in the dataset. This aggregated score quantifies whether cells with different roles in the CCC pathway (senders, receivers, *etc.*) are uniformly distributed across cell types (low score) or clearly divided along cell types (high score). Conversely, the overall heterogeneity score of a cell type is obtained by averaging the heterogeneity scores of cell type *i* over all signaling pathways and informs whether cells of that type assume different communication behaviors across many CCC pathways (low score) or mostly behave consistently (high score).

To define a spatial heterogeneity score, the definition of eq. (3) is modified to sum over all cells within a cutoff distance from a given spatial coordinate (𝑥_*i*_, 𝑦_*i*_).

6. **Statistical analysis of signaling mode robustness.** Given a CCC pathway *P* with *K* associated CCC signaling modes, cells within any cell type will split between these *K* modes (for example: the SPN-3 cell type splits between *K* = 3 signaling modes in the EGF pathway including with sender, sender/receiver, and weak modes – see Fig. 3C). We seek to discriminate between situations where (1) the cell type is largely composed of cells from a single mode, and it is thus possible to average CCC behavior over the entire cell type, or (2) the cell type is a mixture of multiple modes and cell type-level averaging would lead to significant loss of information. Let *N*_*i*_ be the number of cells in cell type *i*, and *N*_*ij*_ the fraction of cells in cell type *i* that were clustered in CCC mode *j*. Let *n*_*ij*_ be a Poisson-distributed random variable with mean and standard deviation equal to 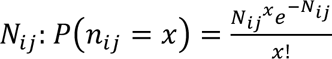. Then, we define a p-value as the probability to randomly sample a number of cells equal or larger than *N*_*i*_. BY default, the threshold value is set to a default value 𝜏 = 0.05 following standard convention for p-value thresholding and based on a sample of 10000 random numbers, but the parameters can be modified by users.

7. **Model of cell-cell communication.** First, a score of the communication via a specific CCC pathway *S*_*ij*_ is defined between each pair of cells in the dataset as a product of signaling intensity (𝐼_*ij*_) and intracellular response (*K*_j_):

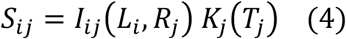

Based on this definition, communication from cell *i* to cell *j* requires both (1) expression of ligands/receptors as well as (2) activation of the pathway’s downstream targets in the receiving cell. First, 𝐼_*ij*_(𝐿_*i*_, 𝑅_j_) is the intensity whereby 𝐿_*i*_ and 𝑅_j_represent the average expression of ligand and receiver in cells *i* and *j*, respectively. ScRICH implements two possible models for signaling intensity, including a diffusion model:

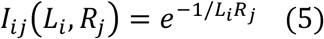

And a Michaelis-Menten model:

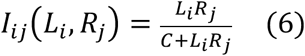

To normalize the variable copy numbers associated to different pathways, the product 𝐿_*i*_𝑅_j_is rescaled within [0,1] by normalizing to the largest value of 𝐿_*i*_𝑅_j_ among all cell pairs in the dataset. Based on this normalization, we set 𝐶 = 0.5 in eq. 6. The user can choose between the diffusion and Michaelis-Menten models, and further comparison is presented in Supplementary Figure S2. Second, *K*_j_(𝑇_j_) quantifies the intracellular response in the receiving cell as:

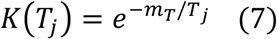

Where 𝑇_j_ is the activation of downstream pathways in cell *j* and 𝑚_𝑇_ is the number of downstream targets. The activation is defined either as the average expression of downstream pathways in the receiving cell:

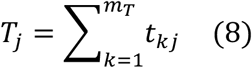

Or, in the “RNA velocity communication model”, as the average RNA velocity of the downstream targets in the receiving cell:

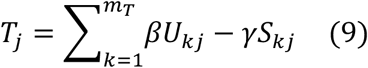

Where 𝑈_*k*j_ and *S*_*k*j_ are the unspliced and spliced copy numbers of the k-th downstream target in cell *j*, and *β* and 𝛾 are the splicing and degradation rate constants computed using the scVelo package^30^.

Finally, the dense cell-to-cell communication network is coarse-grained to a meta-cell to meta-cell network by aggregating the cell-pair communication scores based on the meta-cell assignment that summarizes both cell type and CCC signaling modes that was described in a previous section.

8. **Emerging CCC patterns.** After CCC modes are identified for multiple (K) CCC signaling pathways, each cell in the dataset is described by a K-long vector whereby the i^th^ element represents the cell’s membership to the i^th^ CCC signaling pathway’s mode. The K observables are used for unsupervised clustering using the Leiden algorithm, whereby each cluster represents a CCC pattern.

9. **Pathway redundancy.** Given a system where cells are clustered in N_1_ and N_2_ CCC modes along two CCC signaling pathways (S_1_ and S_2_), let *P*_1_(*j*) and *P*_2_(*k*) be the probabilities to observe a cell in the j-th CCC mode of pathway S_1_ and the k-th mode of pathway S_2_, respectively, and *P*(*j*, *k*) be the joint probability to observe a cell in the j-th CCC mode for pathway S_1_ and the k-th CCC mode for pathway S_2_. The redundancy between the two pathways is defined as:

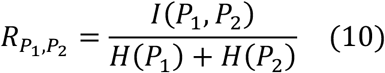

Where 𝐼(*P*_1_, *P*_2_) is the mutual information between *P*_1_ and *P*_2_, and *H*(*P*_1_) and *H*(*P*_2_) are the entropies of *P*_1_ and *P*_2_. With the definition of eq. (10), 𝑅_*P*1,*P*2_ is semi-positive and assumes a minimum value of 0 if and only if the membership to CCC modes for the two pathways are completely uncorrelated.

10. **Inference of gene regulatory network.** To construct the coupled gene regulatory network linking two CCC pathways (i.e., EGF and TGFB), we first compiled a gene set including: (1) the set of ligands and receptor of the two signaling pathways and (2) the set of downstream genes associated with the two pathways. ScRICH includes a built-in database for ligand, receptor, and intracellular mediators, but offers the option to provide user inputs. Then, the GRN is inferred from the scRNA-seq data object including the unspliced and spliced layers and the specified gene set using the spliceJAC package^32^.

11. **Comparison with other CCC inference methods.** The benchmarking against existing CCC inference methods includes 3 steps: (1) Calculation of ScRICH’s CCC signaling matrix; (2) calculation of other method(s)’s CCC signaling matrix; and (3) comparison via multiple scoring metrics. First, ScRICH is run on a given CCC signaling pathways. The resulting CCC network, however, is averaged over the CCC signaling modes, resulting in a CCC matrix between cell types. Second, we compute the same CCC matrix using existing methods using CellChat, CellPhoneDB and Liana+ using their standard pipeline and default settings. Liana+ integrates multiple CCC inference methods to provide a consensus prediction for each distinct ligand-receptor pair^19^. These pairs were aggregated into CCC pathways using the Cellchat dataset (for example, interactions between Jag1-Notch1 and Jag2-Notch2 are aggregated into a unique interaction for Notch pathway). The output CCC matrices from these methods describe interactions between cell types and are thus directly comparable with ScRICH’s coarse-grained output. Finally, the CCC matrices are compared based on 3 scoring metrics. (1) Spearman correlation is computed on the one-dimensional vectors obtained by flattening the two-dimensional CCC matrices. (2) Matrix distance is a squared sum of pairwise distances between matrix elements:

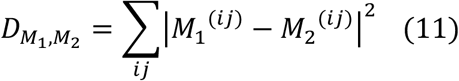

Where the sum runs over the matrix’s rows (*i*) and columns (*j*). (3) Interaction overlap is the fraction of connection between cell types that are identified in both models, regardless of interaction amplitude.

12. **Data preprocessing.** OVCA420 time course data was preprocessed, normalized, regressed and scaled using scanpy’s standard workflow^69^ (min_genes=200, min_cells=3, genes_by_counts<3500, total_counts<1500, target_sum=1e4, max_value=10). Highly variable genes were identified (min_mean=0.0125, max_mean=3, min_disp=0.5), followed by clustering with n_pcs=40, n_neighbors=10. The erythroid differentiation dataset was downloaded and preprocessed using scvelo’s built-in functions after setting min_shared_counts=20. For skin organoid and OVCA420 datasets, RNA velocity was computed following scvelo’s standard workflow with n_pcs=30, n_neighbors=30. For the erythroid differentiation dataset, RNA velocity was computed following Unitvelo’s tutorial on the same dataset maintaining the same parameter values^39^. For the developing ckicken heart spatial transcriptomics dataset, RNA velocity were computed following SIRV’s tutorial on the same dataset maintaining the same parameter values^74^.

13. **FibHSE organotypic culture.** Human primary keratinocytes and dermal fibroblasts were isolated from discarded neonatal foreskin, as described previously^75,76^. Primary human keratinocytes were maintained below 70% confluency in K-SFM medium (Keratinocyte-SFM medium supplemented with 50 μg/ml Bovine Pituitary Extract, 5 ng/ml human recombinant Epidermal Growth Factor 1–53 and 100 U/mL penicillin-streptomycin). Primary fibroblasts were maintained in complete DMEM (DMEM supplemented with 10% fetal bovine serum and 100 U/mL penicillin-streptomycin). Split-thickness human skin was purchased from the New York Firefighters Skin Bank (New York, New York, USA). FibHSEs were generated using a modified protocol as outlined Li and Sen^77^. Briefly, primary fibroblasts (1×10^6^ cells) were seeded onto the reticular side of devitalized de-epidermal human dermis (DED) in 1.5 mL of complete DMEM, followed by centrifugation at 600 g for 1 hour. A suspension of primary keratinocytes (1×10^6^ cells in 100μL) was then seeded onto the papillary side of the DED and cultured at an air-liquid interface (ALI) to promote differentiation and stratification over 14 days, with media changes every other day. To assess the impact of EGF supplementation, an EGF-free culture medium was introduced on day 7 of ALI.

14. **Immunofluorescence staining.** Immunofluorescence (IF) staining were performed as previously described^33^ with slight modifications. In brief, frozen slides containing 20μm organoid sections were fixed with 4% paraformaldehyde (PFA) for 15 min, blocked in PBS containing 2% bovine serum albumin (BSA) and 0.1% Triton X-100 for 1h, and co-stained with either rabbit anti-EGF (1:500; Proteintech; 27141-1-AP) and mouse anti-EGFR (1:500; Proteintech; 66455-1-Ig), or rabbit anti-HRAS (1:500; Proteintech; 18295-1-AP) and mouse anti-INHBA (1:500; Proteintech; 60352-1-Ig). Slides were washed and then incubated with Alexa Fluor 546 Goat Anti-Mouse IgG (H+L) (Invitrogen) and Alexa Fluor 647 Goat Anti-Rabbit IgG (H+L) (Invitrogen). Mounting medium (abcam) containing 4,6-diamidino-2-phenylindole (DAPI) was used to visualize nuclei. All slides were immunostained in a single batch to ensure consistency across samples. Images were captured during the same experimental session using consistent settings on a ZEISS LSM780 confocal microscope at 20x magnification. For each condition, three different fields of tissue sections were acquired from two independent FibHSE replicates. Colocalization analysis was performed using the *Colocalization Finder* plugin in ImageJ.

## Code availability

The code developed to run scRICH is freely available on Github at https://github.com/federicobocci/ScRICH.

## Data availability

All sequencing data analyzed in this project is publicly available through NCBI’s Gene Expression Omnibus (GEO) database with accession numbers GSE190695 (human-skin-equivalent organoids), GSE147405 (time course of OVCA420 cells), GSE149457 (Spatial transcriptomics of chicken hear development). The erythroid differentiation dataset was originally published by Pijuan-Sala et al^78^, but was accesses through scvelo’s built-in *datasets.gastrulation_erythroid()* function^30^.

## Acknowledgement

The work was partly supported by National Science Foundation grants DMS1763272 and CBET2134916, and National Institutes of Health grants R01GM152494 and R01AR079150.

