## Supplementary figures and tables for "Robust identification of cell-cell communication heterogeneity in single cells"

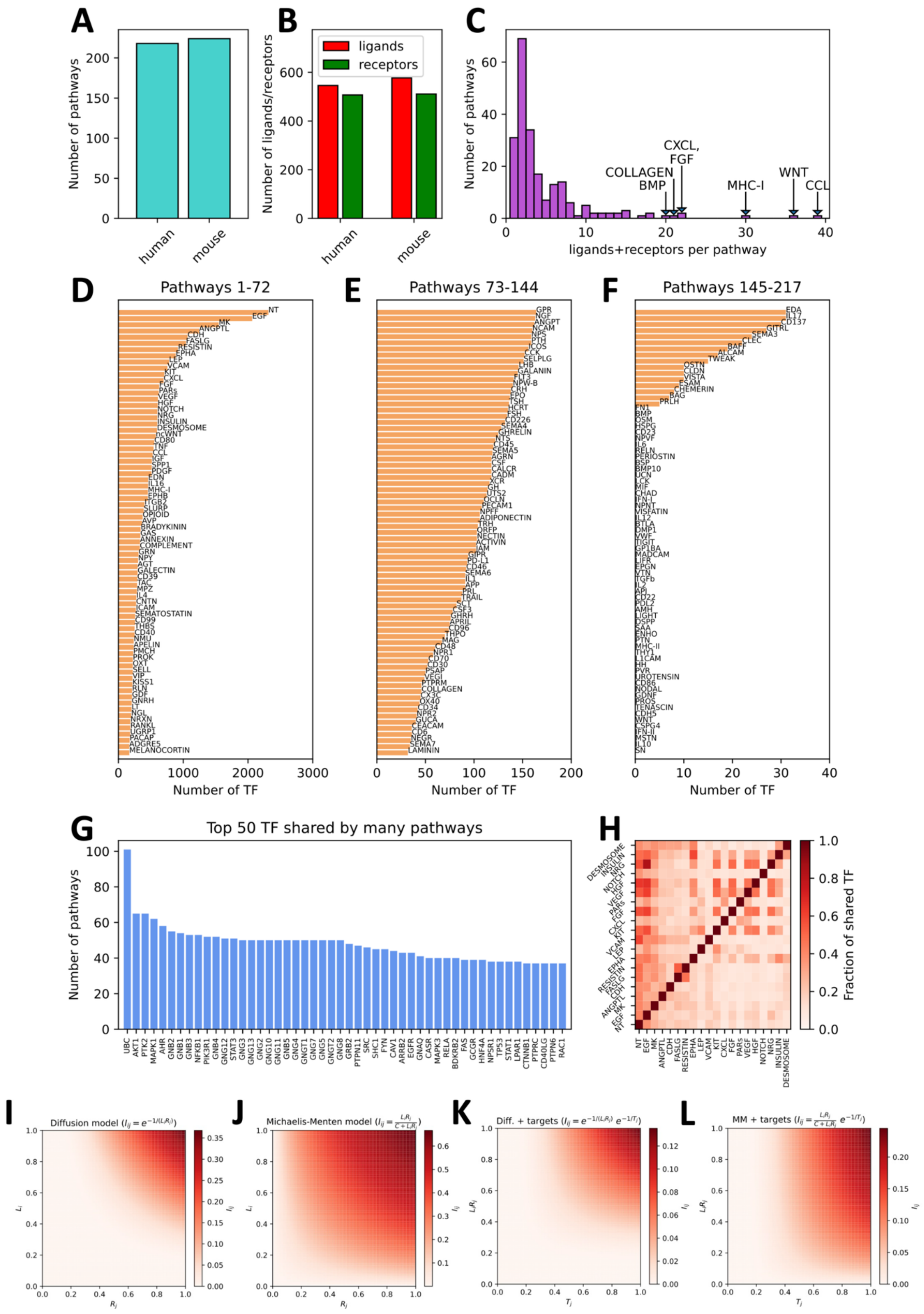
A
Number of pathways
human
mouse
B
Number of ligands/receptors
ligands
receptors
human
mouse
C
Number of pathways
ligands+receptors per pathway
COLLAGEN
BMP
CXCL,
FGF
MHC-I
WNT
CCL
D
Pathways 1-72
Number of TF
NT
EGF
MK
ANGPTL
CDH
FASLG
RESISTIN
EPHA
LEP
VCAM
KIT
CXCL
FGF
PARs
VEGF
HGF
NOTCH
NRG
INSULIN
DESMOSOME
ncWNT
CD80
TNF
CCL
IGF
SPP1
PDGF
EDN
IL16
MHC-I
EPHB
ITGB2
SLURP
OPIOID
AVP
BRADYKININ
GAS
ANNEXIN
COMPLEMENT
GRN
NPY
AGT
GALECTIN
CD39
TAC
MPZ
IL4
CNTN
ICAM
SEMATOSTATIN
CD99
THBS
CD40
NMU
APELIN
PMCH
PROK
OXT
SELL
VIP
KISS1
RLN
GDF
GNRH
LT
NGL
NRXN
RANKL
UGRP1
PACAP
ADGRE5
MELANOCORTIN
E
Pathways 73-144
Number of TF
GPR
NGF
ANGPT
NCAM
NPS
PTH
ICOS
CCK
SELPLG
LHB
GALANIN
FLT3
NPW-B
CRH
EPO
TSH
HCRT
FSH
CD226
SEMA4
GHRELIN
NTS
CD45
SEMA5
AGRN
CSF
CALCR
CADM
XCR
GH
UTS2
OCLN
PECAM1
NPFF
ADIPONECTIN
TRH
QRFP
NECTIN
ACTIVIN
JAM
GIPR
PD-L1
CD46
SEMA6
IL1
APP
PRL
TRAIL
SCT
CSF3
GHRH
APRIL
CD96
THPO
MAG
CD48
NPR1
CD70
CD30
PSAP
VEGI
PTPRM
COLLAGEN
CX3C
OX40
CD34
NPR2
GUCA
CEACAM
CD6
NEGR
SEMA7
LAMININ
F
Pathways 145-217
Number of TF
EDA
IL17
CD137
GITRL
SEMA3
CLEC
BAFF
ALCAM
TWEAK
OSTN
CLDN
VISTA
ESAM
CHEMERIN
BAG
PRLH
FN1
BMP
OSM
HSPG
CD23
NPVF
IL6
RELN
PERIOSTIN
BSP
BMP10
UCN
LCK
MIF
CHAD
IFN-I
NPNT
VISFATIN
IL12
BTLA
DMP1
VWF
TIGIT
GP1BA
MADCAM
LIFR
EPGN
VTN
TGFb
IL2
API
CD22
PDL2
AMH
LIGHT
OSPP
SAA
ENHO
PTN
MHC-II
THY1
L1CAM
HH
PVR
UROTENSIN
CD86
NODAL
GDNF
PROS
TENASCIN
CDH5
WNT
CSPG4
IFN-II
MSTN
IL10
SN
G
Top 50 TF shared by many pathways
Number of pathways
UBC AKT1 PTK2 MAPK1 AHR GNB2 GNB1 GNB3 NFKB1 PIK3R1 GNB4 GNG12 STAT3 GNG3 GNG13 GNG2 GNG10 GNG11 GNB5 GNG4 GNGT1 GNG7 GNG5 GNGT2 GNG8 GRB2 PTPN11 SRC SHC1 FYN CAV1 ARRB2 EGFR GNAQ CASR MAPK3 RELA BDKRB2 FAS GCGR HNF4A NPSR1 TP53 STAT1 LPAR1 CTNNB1 PTPRC CD40LG PTPN6 RAC1
H
Fraction of shared TF
NT EGF MK ANGPTL CDH FASLG RESISTIN EPHA LEP VCAM KIT CXCL FGF PARs VEGF HGF NOTCH NRG INSULIN DESMOSOME
I
Diffusion model ($I_{ij} = e^{-1/(L_iR_j)}$)
$L_i$
$R_j$
$I_{ij}$
J
Michaelis-Menten model ($I_{ij} = \frac{L_iR_j}{C+L_iR_j}$)
$L_i$
$R_j$
$I_{ij}$
K
Diff. + targets ($I_{ij} = e^{-1/(L_iR_j)}\, e^{-1/T_j}$)
$L_iR_j$
$T_j$
$I_{ij}$
L
MM + targets ($I_{ij} = \frac{L_iR_j}{C+L_iR_j}\, e^{-1/T_j}$)
$L_iR_j$
$T_j$
$I_{ij}$

**Supplementary figure 1. Overview of the scRICH database. (A)** Number of cell-cell communication signaling pathways for human and mouse. **(B)** Total number of ligands and receptors for human and mouse. **(C)** Histogram of number of ligands + receptors per pathway. Annotations highlight the pathways with higher number of ligands + receptor. **(D-E-F)** Number of transcription factors (TF) directly connected to each pathway's set of receptors. The pathways are split between the 3 panels for ease of visualization. **(G)** The top 50 TF that are directly connected to the most pathways. **(H)** The TF overlap between the top 20 pathways (based on panel D). The overlap is quantified as the fraction of TF that are shared between the pair of pathways. **(I-L)** Comparison of signaling models between meta-cells. **(I)** The signaling between meta-cells $i$ (the sender) and meta-cell $j$ (the receiver) as a function of receiver and ligand levels in the simplified diffusion model without downstream target. **(J)** The signaling between meta-cells in the simplified Michaelis-Menten model without downstream target. While both models are symmetrical with respect to $L_i$ and $R_j$, the signaling in the Michaelis-Menten model increases gradually, whereas the signaling exhibits a drastic cut-off when $L_i$ and $R_j$ are below threshold values. **(K)** The signaling between meta-cells as a function of receiver-ligand product $L_i R_j$ and downstream target levels in the receiver meta-cell $T_j$ in the diffusion model. **(L)** Same as panel C in the Michaelis-Menten model with downstream target.

<u>Notes:</u> (1) For simplicity, it is assumed here that only one species of ligand, receptor, and target exist. ScRICH averages expression levels in presence of multiple species. (2) In the ScRICH framework, the set of all interactions between meta-cells for a specific cell-cell communication pathway are sum-normalized to 1, thus resolving the different value ranges assumed by the different models. (3) $C = 1$ in both panels J and L, since it is assumed that ligand and receptor expression are re-normalized in [0,1]. (4) $T_j$ here is loosely defined as the "target level"; In ScRICH, $T_j$ can be either quantified by the gene expression or RNA velocity of the target gene(s).

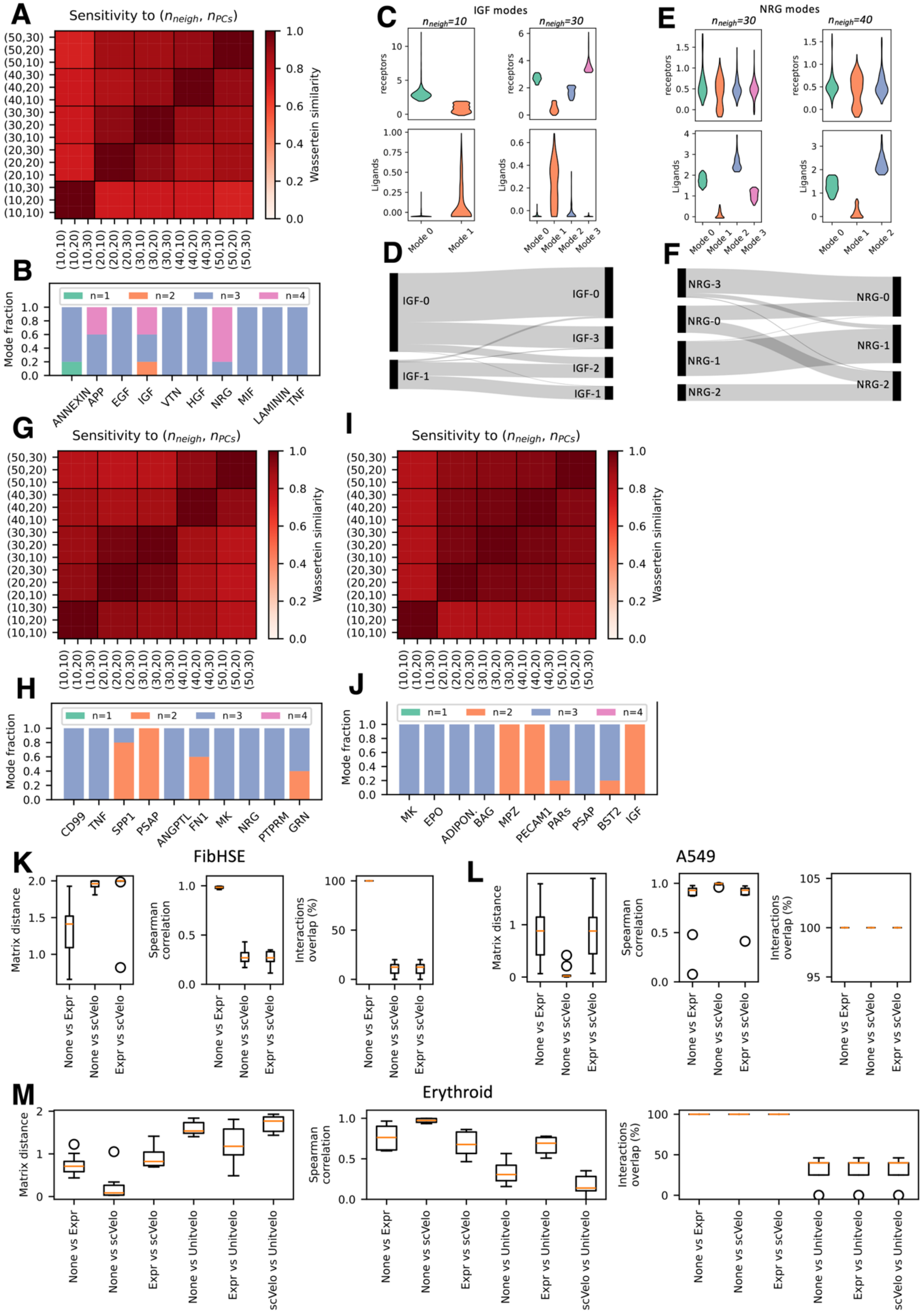
A
Sensitivity to ($n_{neigh}$, $n_{PCs}$)
Wassertein similarity
B
Mode fraction
n=1 n=2 n=3 n=4
ANNEXIN APP EGF IGF VTN HGF NRG MIF LAMININ TNF
C
IGF modes
$n_{neigh}$=10
$n_{neigh}$=30
receptors
Ligands
Mode 0 Mode 1 Mode 2 Mode 3
D
IGF-0 IGF-1 IGF-0 IGF-3 IGF-2 IGF-1
E
NRG modes
$n_{neigh}$=30
$n_{neigh}$=40
F
NRG-3 NRG-0 NRG-1 NRG-2 NRG-0 NRG-1 NRG-2
G
Sensitivity to ($n_{neigh}$, $n_{PCs}$)
H
CD99 TNF SPP1 PSAP ANGPTL FN1 MK NRG PTPRM GRN
I
Sensitivity to ($n_{neigh}$, $n_{PCs}$)
J
MK EPO ADIPON. BAG MPZ PECAM1 PARs PSAP BST2 IGF
K
FibHSE
Matrix distance
Spearman correlation
Interactions overlap (%)
None vs Expr
None vs scVelo
Expr vs scVelo
L
A549
M
Erythroid
None vs Unitvelo
Expr vs Unitvelo
scVelo vs Unitvelo

**Supplementary figure 2. Benchmarking and parameter testing. (A)** Similarity based on Wassertein distance of cell-cell communication modes for the top 10 pathways identified in the FibHSE dataset for different combinations of number of nearest neighbors and principal components. **(B)** Fraction of mode clustering solutions with different number of modes across all parameter combinations from (A) for each CCC pathway. **(C)** CCC modes for the IGF pathway for two parameter combinations yielding different number of modes. **(D)** Mapping of IGF signaling modes for the two parameter combinations. **(E-F)** Same as (C-D) for the NRG pathway. **(G-J)** Same as (A-B) for the A549 dataset analyzed in Fig. 6 (G-H) and for the erythroid differentiation dataset analyzed in Fig. 5 (I-J). **(K)** Comparison of ScRICH CCC communication prediction in FibHSE for the top 10 CCC pathways based on standalone ligand/receptor expression (None), ligand/receptor and target gene expression (Expr), and (3) ligand/receptor expression plus RNA velocity of target genes (scVelo). Comparison of CCC interaction matrix is based on matrix distance, Spearman correlation, and overlap similar to Fig. 2H. **(L)** Same as (K) for the A549 dataset. **(M)** Same as (K) for the erythroid differentiation dataset including a comparison between scVelo-based and Unitvelo-based RNA velocity inference.

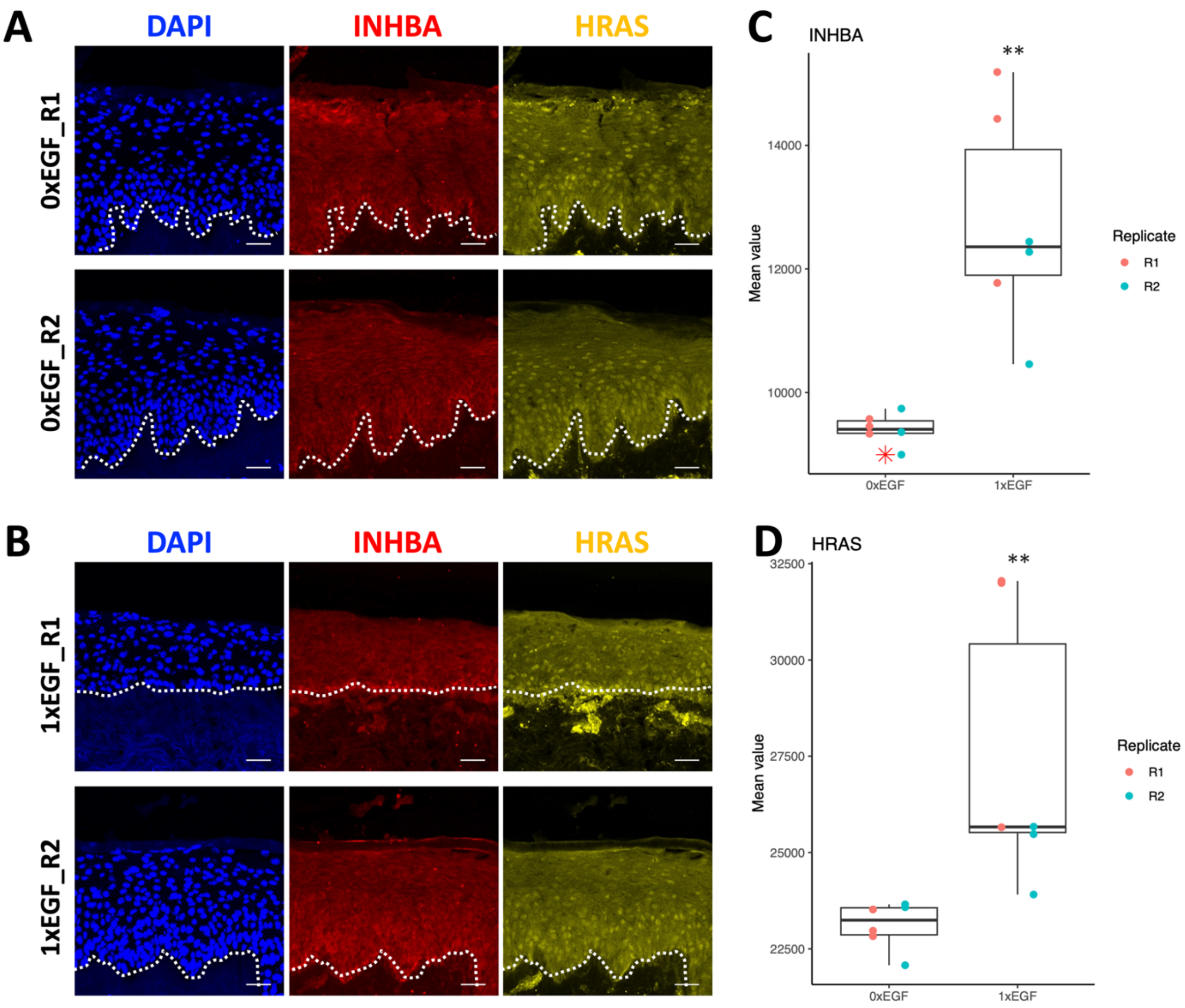

**Supplementary Figure S3. Expression of HRAS and INHBA in FibHSE. (A-B)** Co-localization of HRAS and INHBA in FibHSE for the cases of no exogenous EGF (A, 0XEGF) and soluble EGF in solution (B, 1XEGF). R1 and R2 indicate replicates for each condition. **(C-D)** Quantification of INHBA expression based on replicates in C and EGF dosage. **(D)** Same as (C) for HRAS.

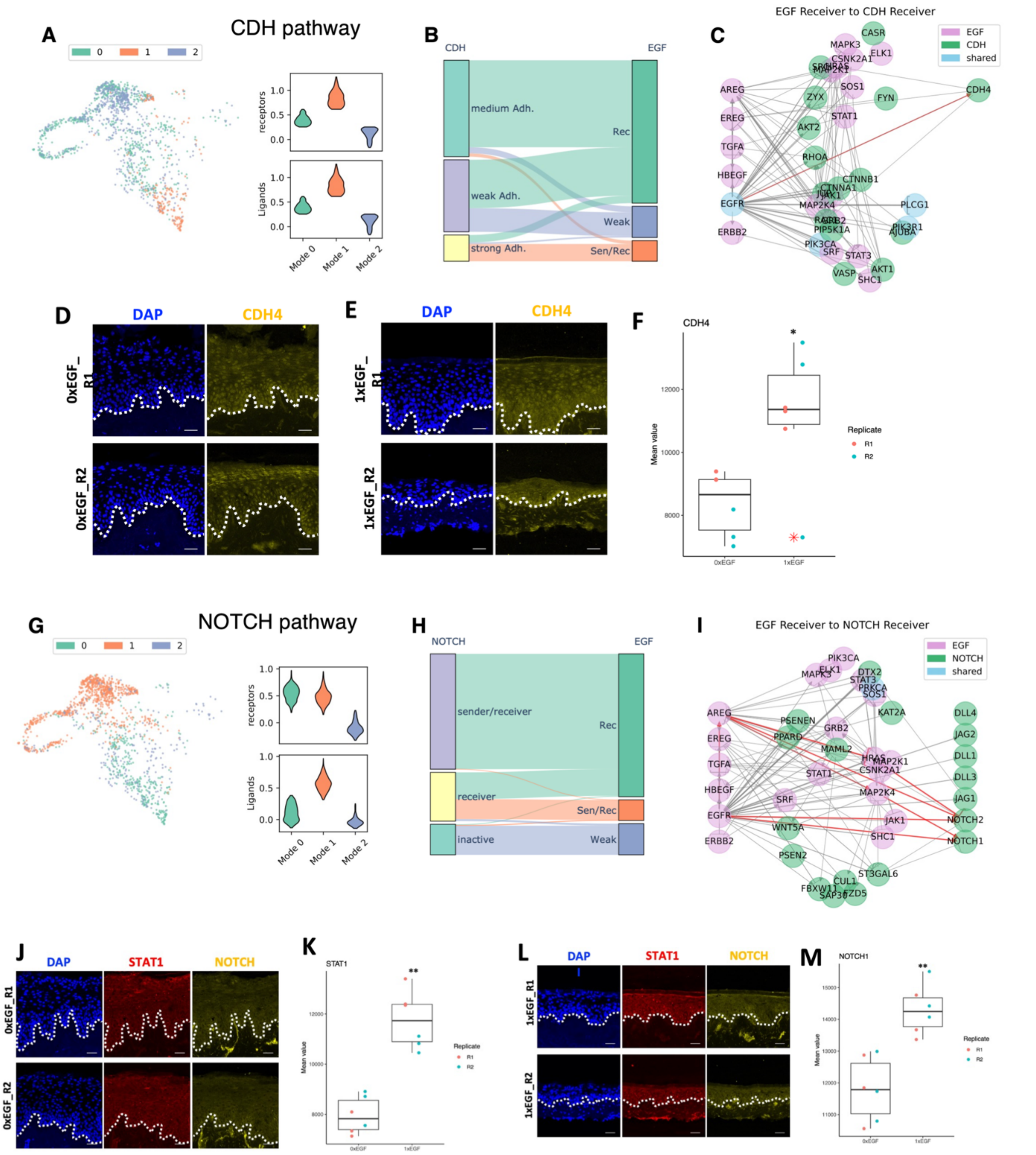

**Supplementary figure S4. Connection between EGF and other EMT-related CCC pathways. (A)** The CDH CCC modes including low, medium and high adhesion. **(B)** Sankey diagram of correspondence between CDH and EGF CCC modes. **(C)** The gene regulatory network connecting the EGF and CDH pathways in the EGF receiver cells. **(D-E)** Co-localization of CDH4 in FibHSE for the cases of no exogenous EGF (0XEGF) and soluble EGF in solution (1XEGF). R1 and R2 indicate replicates for each condition. **(F)** Quantification of CDH4 expression based in replicate and EGF dosage. **(G-H-I)** Same as A-B-C for the Notch

CCC pathway. **(J, L)** Co-localization of STAT1 and NOTCH in FibHSE for the cases of no exogenous EGF (0XEGF) and soluble EGF in solution (1XEGF). R1 and R2 indicate replicates for each condition. **(K, M)** Quantification of STAT1 and NOTCH expression based in replicate and EGF dosage.

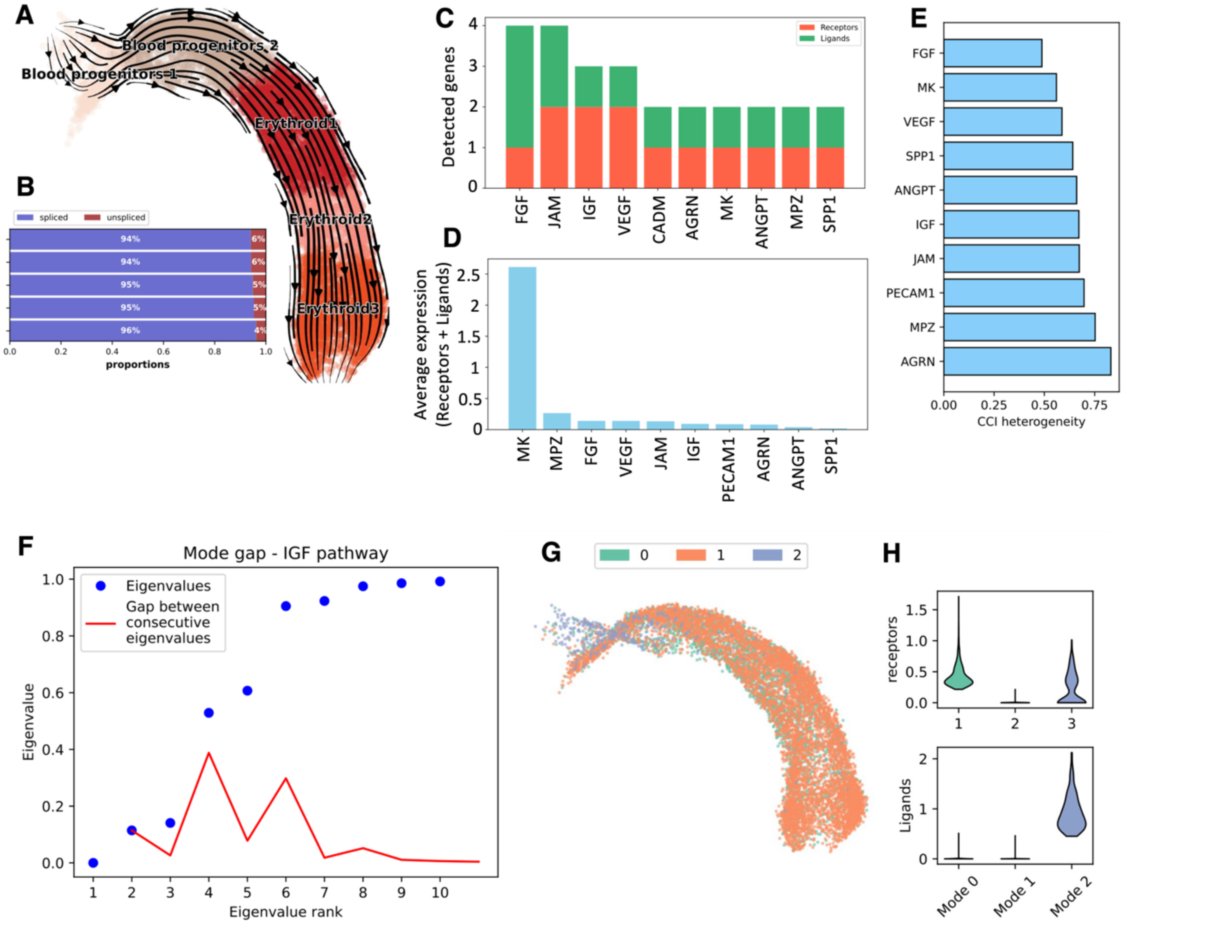

**Supplementary figure S5. Supplementary analysis of erythroid cell differentiation. (A)** UMAP, cell types, and RNA velocity of erythroid cell lineage. **(B)** Fraction of unspliced RNA counts in each cell type. **(C-D)** The top CCC pathways identified based on total ligands and receptors expressed (top) and ligand/receptor expression (bottom). **(E)** CCI heterogeneity of the top CCC pathway. **(F)** Eigenvalue gap for the IFG CCC pathway highlighting three CCC modes. **(G)** UMAP embedding of the IGF CCC modes. **(H)** Expression of IGF ligands and receptors in the three IFG modes via violinplot. The three modes correspond to receiver, inactive, and sender.

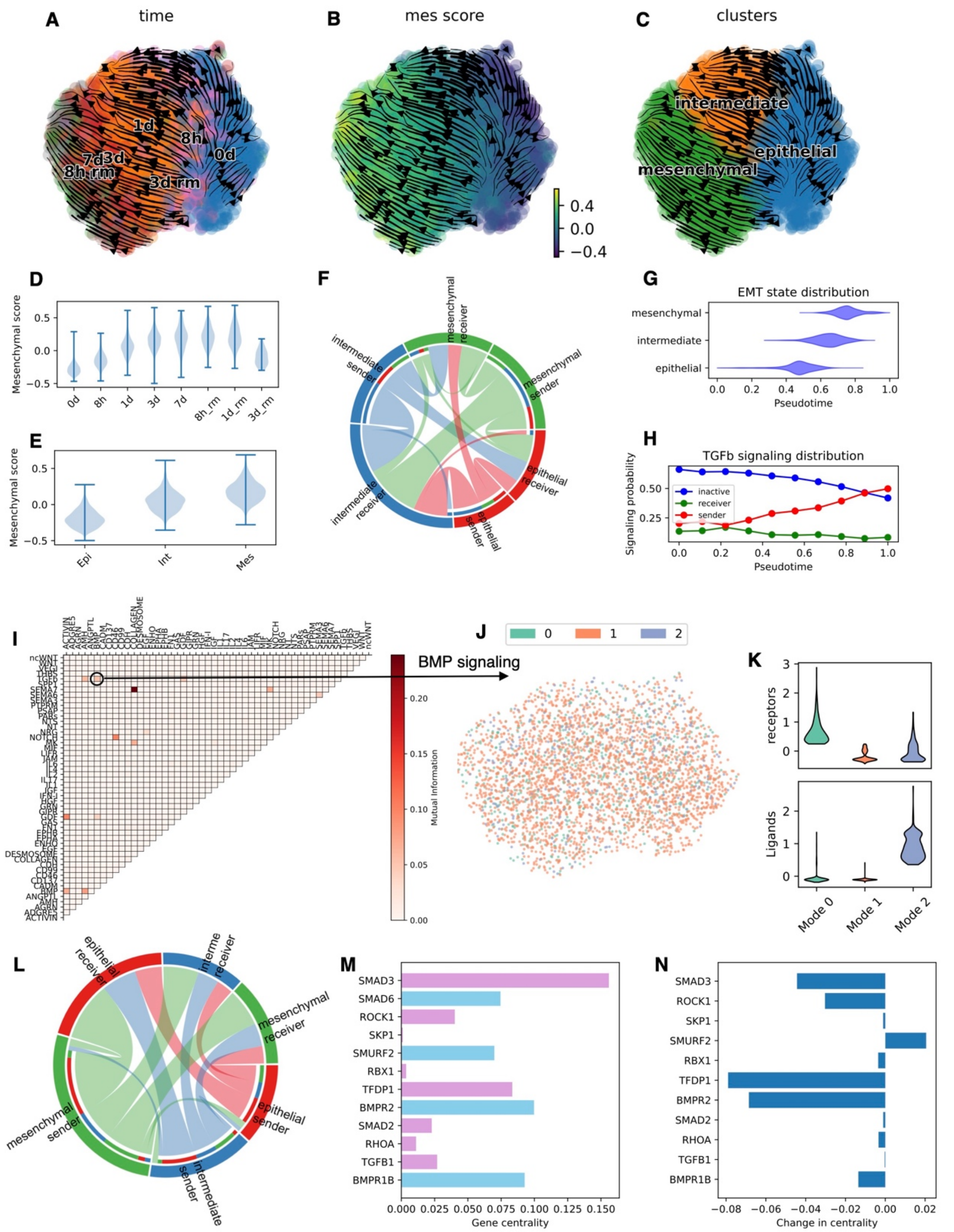

**Supplementary Figure 6. Supplementary analysis of CCC during EMT. (A-B-C)** Low dimensional UMAP embedding and RNA velocity where cells are color-coded by time point (A), mesenchymal score (B), and EMT cluster annotation (C). **(D-E)** The mesenchymal score by time point (D) and cluster (E). **(F)** The TGFb CCC network. **(G)** EMT cell state distribution as a function of pseudotime. **(H)** Fraction of cells in the different TGFb communication modes as a function of pseudotime. **(I)** Pairwise mutual information between CCC pathways. **(J)** The CCC modes of the BMP pathway in UMAP embedding. (K) Expression of BMP receptors and ligands highlighting a BMP receiver (mode 0), inactive (mode 1) and sender (mode 2). **(L)** The BMP CCC network. **(M)** Betweenness centrality of genes in the joint TGFb-BMP gene regulatory network. Purple and blue bars indicate genes that belong to the TGFb module and shared between the two pathways, respectively, following the same notation of Fig. 5H. **(N)** Change in gene betweenness centrality when comparing the TGFb-BMP GRN inferred at t=0 days and t=7 days.

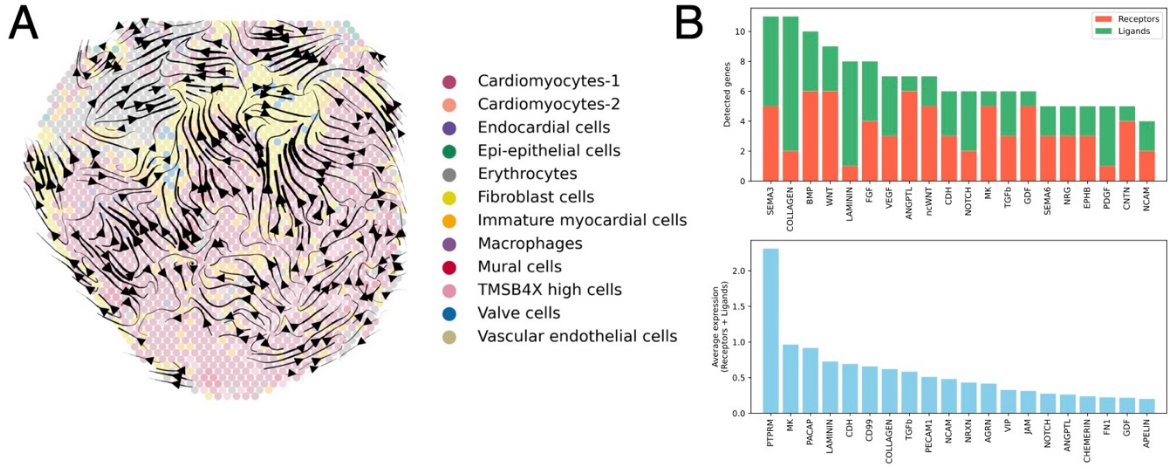

**Supplementary figure S7. Supplementary analysis of chicken earth spatial transcriptomics data. (A)** The cell types and RNA velocity in two-dimensional physical space. **(B)** The top CCC pathways identified based on total ligands and receptors expressed (top) and ligand/receptor expression (bottom).

**Supplementary Tables**

| | |
|---|---|
| EGF | CSNK2A1, EGF, EGFR, ELK1, FOS, GRB2, HRAS, JAK1, JUN, MAP2K1, MAP2K4, MAP3K1, MAPK3, MAPK8, PIK3CA, PIK3R1, PLCG1, PRKCA, PRKCB, RAF1, RASA1, SHC1, SOS1, SRF, STAT1, STAT3, STAT5A |
| TGFB | ACVR1, ACVR1C, ACVR2A, ACVR2B, ACVRL1, AMH, AMHR2, BMP2, BMP4, BMP5, BMP6, BMP7, BMP8A, BMP8B, BMPR1A, BMPR1B, BMPR2, CDKN2B, CHRD, COMP, CREBBP, CUL1, DCN, E2F4, E2F5, EP300, FST, GDF5, GDF6, GDF7, ID1, ID2, ID3, ID4, IFNG, INHBA, INHBB, INHBC, INHBE, LEFTY1, LEFTY2, LTBP1, MAPK1, MAPK3, MYC, NODAL, NOG, PITX2, PPP2CA, PPP2CB, PPP2R1A, PPP2R1B, RBL1, RBL2, RBX1, RHOA, ROCK1, ROCK2, RPS6KB1, RPS6KB2, SKP1, SKP1P2, SMAD1, SMAD2, SMAD3, SMAD4, SMAD5, SMAD6, SMAD7, SMAD9, SMURF1, SMURF2, SP1, TFDP1, TGFB1, TGFB2, TGFB3, TGFBR1, TGFBR2, THBS1, THBS2, THBS3, THBS4, TNF, ZFYVE16, ZFYVE9 |

Supplementary Table T1. The gene sets of CCC pathways obtained from the GSEA database used to infer gene regulatory networks.
